# Extensive and Persistent Extravascular Dermal Fibrin Deposition Characterizes Systemic Sclerosis

**DOI:** 10.1101/2023.01.16.523256

**Authors:** Jeffrey L. Browning, Jag Bhawan, Anna Tseng, Nicholas Crossland, Andreea M Bujor, Katerina Akassoglou, Shervin Assassi, Brian Skaug, Jonathan Ho

**Affiliations:** Departments of Microbiology, Medicine, Boston University Chobanian & Avedesian School of Medicine, Boston, MA; Departments of Rheumatology, Medicine, Boston University Chobanian & Avedesian School of Medicine, Boston, MA; Departments of Dermatopathology, Medicine, Boston University Chobanian & Avedesian School of Medicine, Boston, MA; Departments of Pathology and Laboratory Medicine, Boston University Chobanian & Avedesian School of Medicine, Boston, MA; National Emerging Infectious Diseases Laboratories, Boston University, Boston, MA; Gladstone Institute of Neurological Disease San Francisco California USA; Department of Neurology, Weill Institute for Neurosciences, University of California San Francisco, San Francisco, CA; Division of Rheumatology, University of Texas Health Science Center, Houston, TX; Section Dermatology University of the West Indies, Mona Jamaica

## Abstract

Systemic sclerosis (SSc) is an autoimmune disease characterized by progressive multiorgan fibrosis. While the cause of SSc remains unknown, a perturbed vasculature is considered a critical early step in the pathogenesis. Using fibrinogen as a marker of vascular leakage, we found extensive extravascular fibrinogen deposition in the dermis of both limited and diffuse systemic sclerosis disease, and it was present in both early and late-stage patients. Based on a timed series of excision wounds, retention on the fibrin deposit of the splice variant domain, fibrinogen α_E_C, indicated a recent event, while fibrin networks lacking the α_E_C domain were older. Application of this timing tool to SSc revealed considerable heterogeneity in α_E_C domain distribution providing unique insight into disease activity. Intriguingly, the fibrinogen-α_E_C domain also accumulated in macrophages. These observations indicate that systemic sclerosis is characterized by ongoing vascular leakage resulting in extensive interstitial fibrin deposition that is either continually replenished and/or there is impaired fibrin clearance. Unresolved fibrin deposition might then incite chronic tissue remodeling.

## Introduction

Systemic sclerosis (SSc) is a rare and serious disease characterized by fibrosis of multiple organs (1–4). Primarily due to the presence of autoantibodies, genetic variants common to a number to other systemic autoimmune diseases and the efficacy of autologous stem cell transplantation, SSc is considered an immunological disease, but the underlying disease drivers(s) remain obscure. The disease occurs in two major forms, a diffuse form (dcSSc) affecting all skin regions and a limited form (lcSSc) where skin lesions are constrained to the extremities and face. Morphea or localized scleroderma is a related skin condition wherein well-demarcated skin lesions occur in any distribution, yet systemic disease is lacking. Early observable events in SSc include Raynaud’s phenomenon, edema, inflammatory infiltrates and capillary changes with some form of vasculopathy widely considered to be an initiating event (5–8). The dermal small vessels undergo intimal hyperplasia with the endothelial cells losing characteristic Weibel-Palade bodies, vascular endothelial-cadherin and endothelial nitric oxide synthetase. These changes eventually culminate in a hypovascular tissue with early capillary loss (8). The potential drivers of vascular damage include T cell attack, autoantibody binding, viral infection, FLI1-insufficiency and pericyte detachment (4, 9). As in any healing wound, vascular changes trigger tissue repair mechanisms leading to fibrosis, but in SSc, the connectivity and balance between autoimmunity, vasculopathy and ensuing organ fibrosis is unclear.

Beyond the observation of “puffy” hands and fingers, vascular leakage has not been studied in SSc. We sought to investigate the extent of vascular leakage into the dermal interstitial space in SSc and determine whether leakage is an initial event or is it a pervasive at all stages. Visualization of extravascular fibrinogen or fibrin deposits in tissue biopsies is a critical method to assess vascular leakage. With intravascular activation of the clotting cascade, fibrinogen undergoes proteolytic cleavage by thrombin initiating polymerization, clot formation and homeostasis (10). Extravascular activation of the cascade commences with fibrinogen entering the interstitial spaces where it is converted to a fibrin meshwork. These processes are carefully regulated by evolutionarily ancient proteolytic cascades that control coagulation and fibrinolysis. Fibrin deposition in the extravascular compartment is a universal aspect of tissue injury where it is a potent driver of inflammation by activating macrophages, microglia and neutrophils via the αMβ2 integrin receptor (11–15). The coagulation system is involved in lung and renal fibrosis and glial scarring in model systems with the involvement of fibrinogen itself or associated elements such as thrombin, PAR-1 and the components of fibrinolysis (16–21). Fibrinogen leakage into the extracellular compartment is especially well-studied in the human CNS in neurological diseases such as multiple sclerosis, traumatic brain injury, stroke, and neurodegenerative diseases (22– 25). Based upon multiple routes of investigation, extravascular fibrin formation is a component of the multiple sclerosis lesion and is associated with inflammation, axonal damage and myelin loss (23, 26). Likewise, tumors are often associated with leaky vasculature where increased extravascular fibrinogen deposition plays a role in matrix maturation and cellular infiltration (27, 28). Given this background, we speculated that vascular leakage of fibrinogen could drive fibrosis in SSc.

Over 60 years ago, Fennel and colleagues reported on the presence of fibrin both in intravascular and extravascular locations in the diseased kidney in SSc boldly declaring that excessive amounts of fibrin are what characterizes SSc (29). Fibrin has also been observed in the diseased lungs of SSc patients (30). Notably, blood-based analyses looking at fibrinogen half-life, various components of fibrinolysis such as D-dimer levels as well as thrombin generation potential have been documented as abnormal in SSc (31–35). Additionally, inhibitors of fibrinolysis, plasminogen activator inhibitor (PAI-1/SerpinE1), protease nexin-1 (PN1/SerpinE2) and α-2-antiplasmin are elevated in SSc and have been linked to fibrosis (36–40). Recently, altered clotting parameters were found to track with the formation of digital cutaneous ulcers (41). These observations are generally consistent with a shift towards a hypercoagulable and a hypofibrinolytic state in SSc (16, 42–44). Considering the improved understanding of the potential roles of fibrin deposition in other disease settings and the lack of analysis of extravascular vascular leakage in SSc, we examined fibrin deposition and found extensive and persistent cutaneous extravascular fibrin deposits in both the early and late stages in SSc.

## Results

### Detection of extensive fibrin deposition in SSc skin biopsies

Extracellular fibrinogen is composed of three protein chains, α, β and γ and we used a well-characterized monoclonal antibody to the fibrinogen-γ (Fib-γ) chain to determine the presence of fibrin deposition in healthy control and SSc skin biopsies in the Boston University SSc repository. We will refer to the observed fibrinogen staining as fibrin deposition (27, 45, 46). In this study, we accessed formalin fixed paraffin embedded tissues; however, in dermatology, direct immunofluorescence with frozen sections is more commonly used to define fibrin deposition. To validate our staining, we utilized lichen planus biopsies where fibrin deposition is observed specifically in the dermo epidermal junction and the superficial dermis (Figure 1A) (47). As expected, fibrin deposition was readily seen bordering the leukocytic band in these sections and deposition was largely absent within the inflamed infiltrated regions. CD34 expression identifies endothelial cells, quiescent fibroblasts, the peri-eccrine stromal layer, some perineural sheath cells and hair follicle adnexal structures. In SSc skin, a dramatic transition occurs where the dermal fibroblasts lose these CD34 antibody epitopes (48–50). Fibrin deposition was detected outside the CD34+ blood vessels (Figure 1A). In this image, CD90 staining reveals pericytes, perivascular adventitial fibroblasts and a CD90+ stromal network in the infiltrated region (51). Using either immunofluorescence or conventional imaging, we found extensive fibrin deposition in both dcSSc and lcSSc SSc skin biopsies (Figure 1B,C). For our routine tissue surveys, Fib-γ was combined with CD34 using conventional IHC where this combination of markers explores both the vascular leakage and fibroblast alterations.

**Figure 1:**
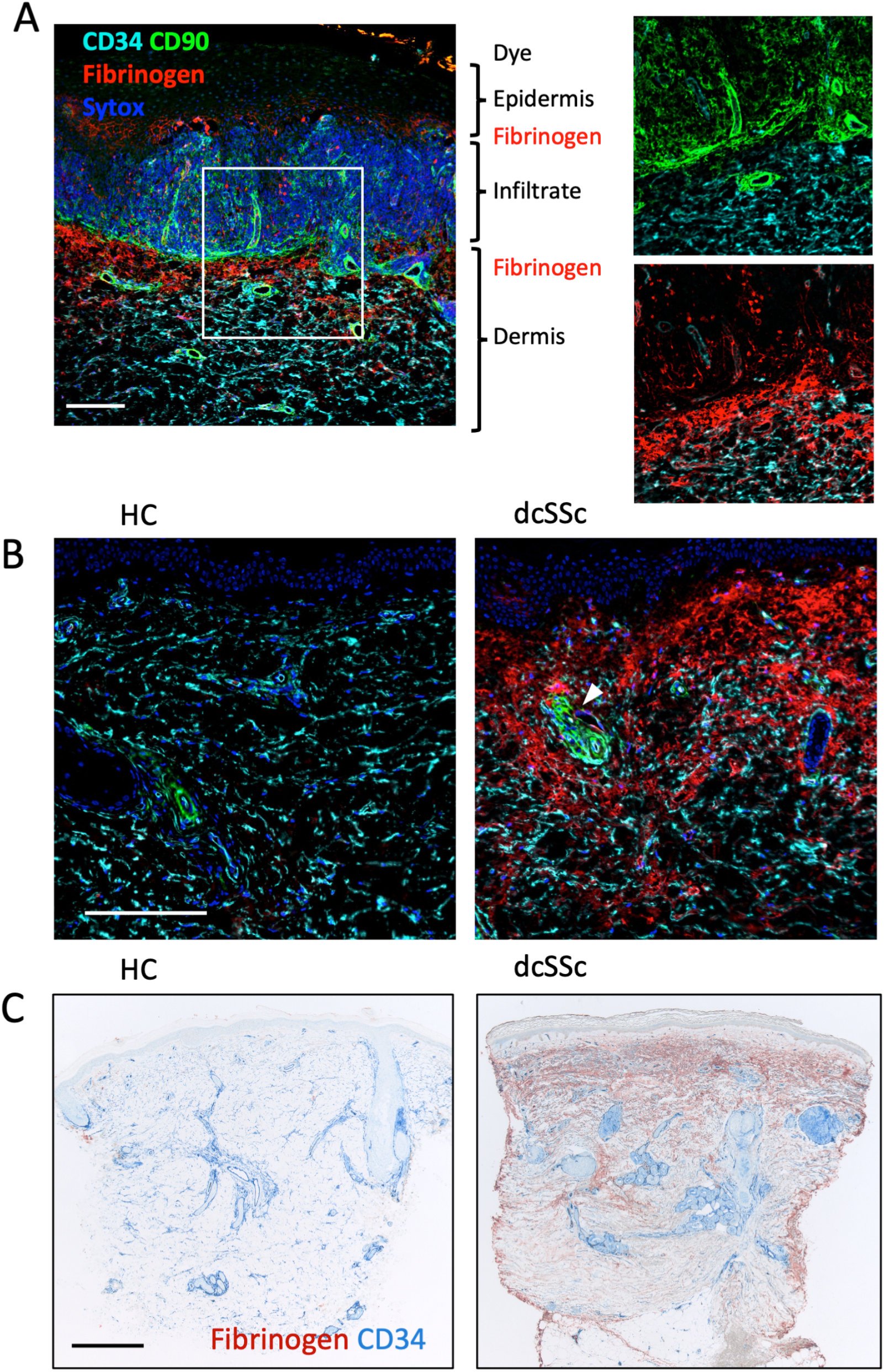
Examples of fibrinogen immunoreactivity in healthy control (HC), lichen planus and SSc skin biopsies. A) Lichen planus is a positive control for fibrinogen staining illustrated here using immunofluorescence imaging. Image shows a heavily infiltrated dermo epidermal interface region with fibrin deposits on both sides of the infiltrate. Inserts reveal the CD90+ reticular network (in the infiltrate), pericytes and perivascular adventitial fibroblasts along with CD34+ vessels and dermal fibroblasts. B) Comparison by immunofluorescence of biopsies from a HC and a dcSSc patient of 8-month duration and a modified Rodnan skin score (mRSS) score of 46. Arrow shows a perivascular adventitial region lacking fibrin deposition. C) Comparison using conventional IHC methods of a HC and a dcSSc patient of 12-year duration with a mRSS of 39. This example is representative of patients with full thickness fibrin deposition. Scale bars are 200 (A,B) and 500 µm (C).

A second anti-human fibrinogen rabbit polyclonal antibody validated the Fib-γ staining patterns (Supplemental Figure 1). The assessment of fibrin deposition in skin punch biopsies is potentially confounded by several factors. Local trauma accompanies the core biopsy and indeed fibrinogen was often seen along the margins of many biopsies including the outer epidermis on occasion (Figure 1C). Therefore, we excluded any trauma-induced margin staining, i.e., fibrin within 0.2 mm of the biopsy margins as well as any narrow waisted sections or shave biopsies from our analysis. In practice these procedure-related regions were easy to identify. Regions containing extravascular erythrocytes were further evidence of procedure-related artifacts and these regions were also excluded. Other factors could potentially influence fibrin deposition such as age, sun exposure damage, and other alterations linked to SSc disease. Sunburn can lead to fibrin deposition and robust leakage is seen in polymorphous light eruption skin, which we also observed (not shown) (52, 53). It is possible that damage due to chronic sun exposure could contribute to fibrin deposition in the upper dermis, however, examination of elastosis in the PRESS biopsy samples (introduced below) showed controls were well matched to SSc patients. In theory, sclerotic skin may require more force to core out the biopsy increasing the extent of the trauma. This possibility is considered unlikely as both 4 mm and 3 mm core biopsies exhibited similar fibrin deposition even though less trauma should occur in the middle of the wider 4 mm biopsies. Roughly 40-50% of SSc patients experience pruritus especially during early phases yet, in an examination of the PRESS samples only one SSc sample exhibited epithelial changes (epidermal lichenification, excoriation or scale formation) possibly consistent with scratching (54). Therefore, scratching is unlikely to be a widespread confounder.

All three fibrinogen genes are highly expressed in the liver, however, Fib-γ is also expressed intracellularly in extrahepatic tissues. To assess whether fibrinogen expression could contribute to the immunohistochemical results, bulk RNA expression databases were analyzed. Identical low levels of Fib-γ RNA were detected in the skin in both control and SSc skin in two independent bulk RNA datasets excluding local gene induction as a contribution to the histological changes (data not shown).

### Fibrin deposition is extravascular in SSc

Some normal skin samples contained small patches of fibrin deposition outside the boundary of CD34+ blood vessels (Figure 2A). Dvorak and colleagues showed that acute fibrinogen leakage into the extravascular space can be detected in normal tissue following injection of agents that increase vascular permeability such as histamine or bradykinin (45). We suspect that transient leaks may occur routinely due to micro-trauma or local mast cell activation creating small fibrin deposits. In SSc, the extensive deposition exhibited multiple patterns including distinct “fibrin deposits with a tangled-like appearance, granular patches and uniform diffuse smears throughout the upper dermis (Figure 2B). The appearance of tangle-like structures in SSc skin resembles deposits described in multiple sclerosis, some tumors and polymorphous light eruption (27, 53, 55). Elastosis was present in some samples with diffuse fibrin, although not all samples with elastosis exhibited diffuse fibrin staining. Extravascular fibrin resided in the interstitial spaces outside the collagen and elastin fibrils (as visualized by elastin autofluorescence or phase contrast, not shown). Fibrin did not co-localize with CD34 or CD90 positive fibroblasts. Intravascular lumen fibrin was observed in some samples, and we could not determine whether this was a consequence of SSc disease or clotting post biopsy excision. In SSc, fibrin distribution was extensive and generally not localized in a defined perivascular manner. However, in some rare SSc biopsies, clear perivascular fibrin deposits were seen (Figure 2C) and, in these cases, it was striking that not all vessels displayed perivascular fibrin suggesting selective leakage. Many SSc biopsies also had extensive fibrinogen deposition in the vicinity of small vessels in peri-eccrine and adipose regions (e.g., Supplemental Figure 1). We propose that vascular leak in SSc leads to extravascular fibrinogen extravasation and fibrin deposition with rapid interstitial dispersion. If most patients have persistent and disseminated fibrin, then a localized perivascular leak may be infrequently captured by biopsy and IHC detection. Three-dimensional imaging of fibrin in large tissue volumes may be used for the detection of localized increases of vascular permeability, as previously shown in the human brain (56).

**Figure 2:**
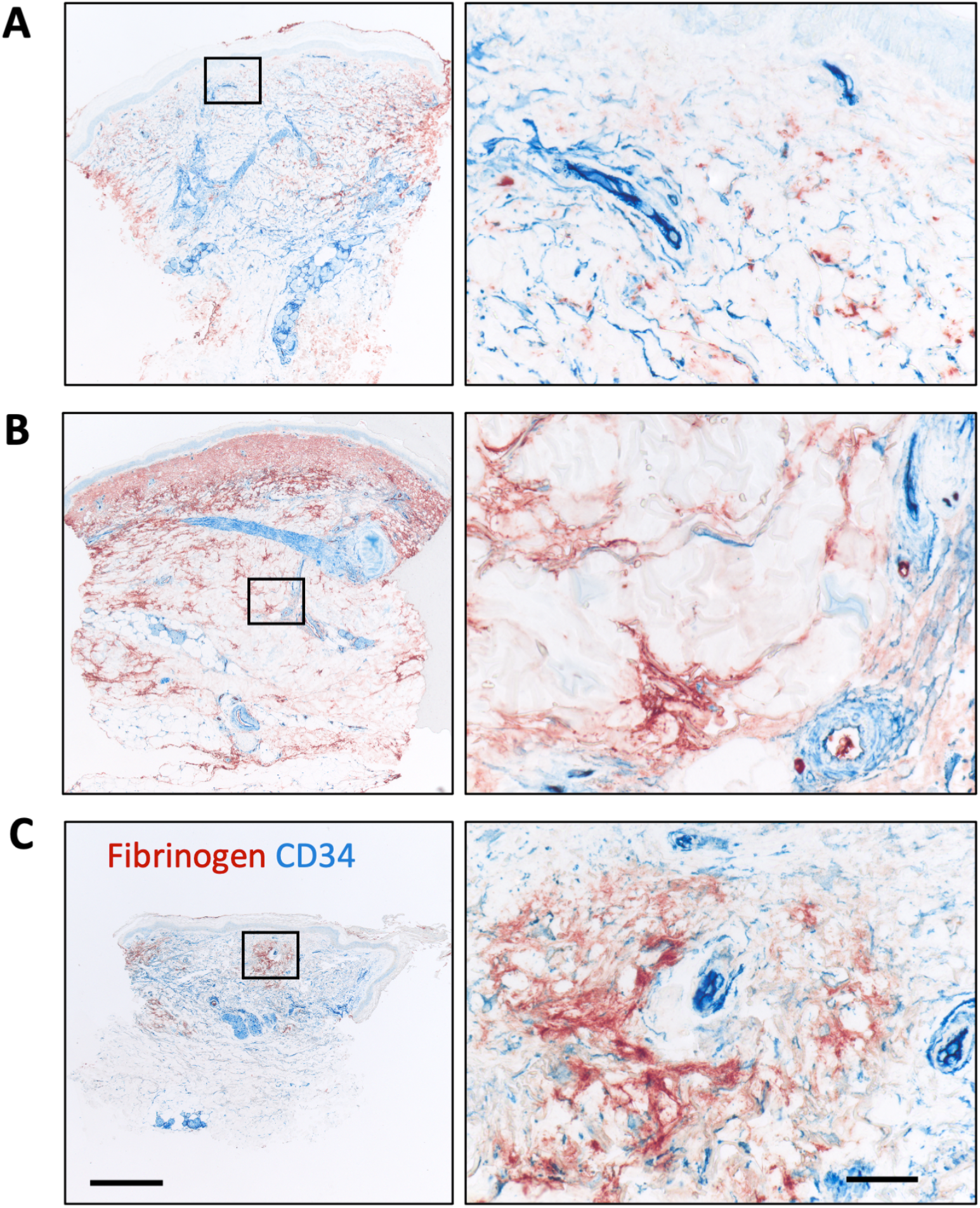
The spectrum of cutaneous fibrin deposition. A) An example of local fibrin deposits in HC skin. B) An example of fibrin “tangle-like” structures seen in SSc along with strong diffuse staining in the upper dermis. C) Rare perivascular fibrin deposits seen in SSc. Scale bars is 500 µm.

### Fibrin deposition is observed in both early and late stages of SSc

Using the Fib-γ mAb and conventional IHC, we scored fibrin deposition on a 0-3 scale and examples of these scores are provided (Supplemental Figure 2). Results for healthy control, dcSSc, lcSSc and morphea are shown in Table I. Elevated fibrinogen scores were seen in both dcSSc and lcSSc, while morphea did not reach statistical significance likely due to low numbers. DcSSc disease progresses over the first 1-2 years and then typically plateaus followed by stabilization in an atrophic state or even with clinical improvement (57–59). In lcSSc, skin involvement is less pronounced, but can also undergo amelioration. Disease duration is generally defined operationally as the time from the first appearance of a non-Raynaud symptom. To determine if fibrin deposition occurs in both the early and late phases of SSc, we exploited both our internal BU repository biopsies as well as a well-characterized PRESS cohort of early disease SSc skin biopsies (median time to diagnosis 1.3 years) (60). We created early (0-2), mid (2-6) and late (>6 years) dcSSc categories for the BU registry. Both PRESS and BU early disease biopsies contained extensive fibrin deposits indicating that deposition is an early event (Table I). Images of all the PRESS cohort biopsies are provided in Supplemental Figures 3-4. Morphometric quantitation of the fibrin positive areas in the PRESS biopsy slides confirmed the visual impression of increased fibrin deposition in SSc as well as the loss of CD34 and a slight increase in macrophage staining (Factor XIIIa+) (Figure 3). Both the mean fibrin scores and the percentage of fibrin+ (e.g., 2+ and 3+) biopsies did not change with disease duration consistent with either continuing extravascular fibrin deposition, impaired clearance, or both. Likewise, both early and late lcSSc skin exhibited extensive fibrin deposition, but our sample size was small.

**Figure 3:**
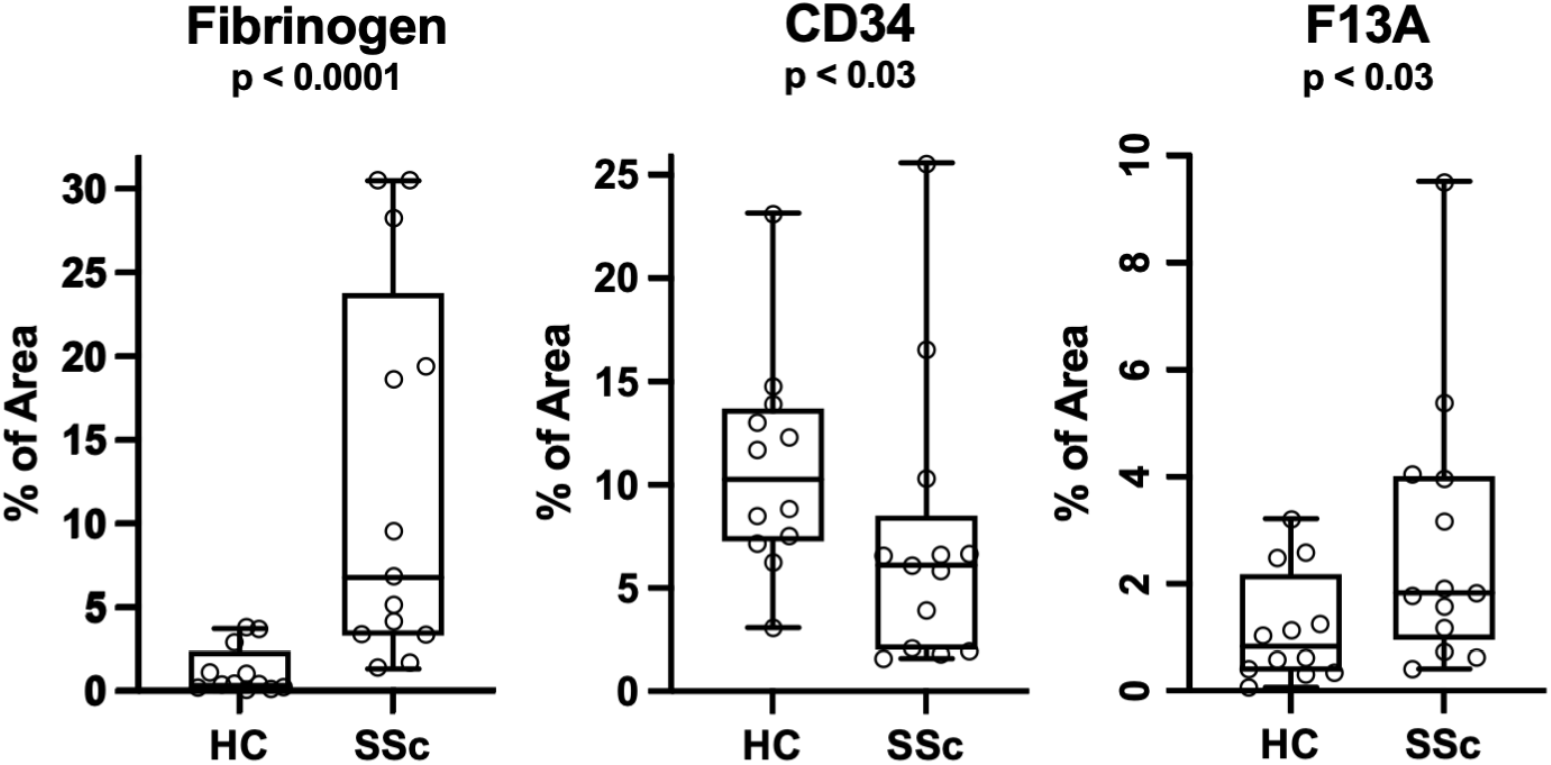
Morphometric analyses of fibrin deposition (A) and CD34 staining (B) in the PRESS cohort of early dcSSc. Significance determined with a Mann-Whitney U test, n = 13.

**Figure 4:**
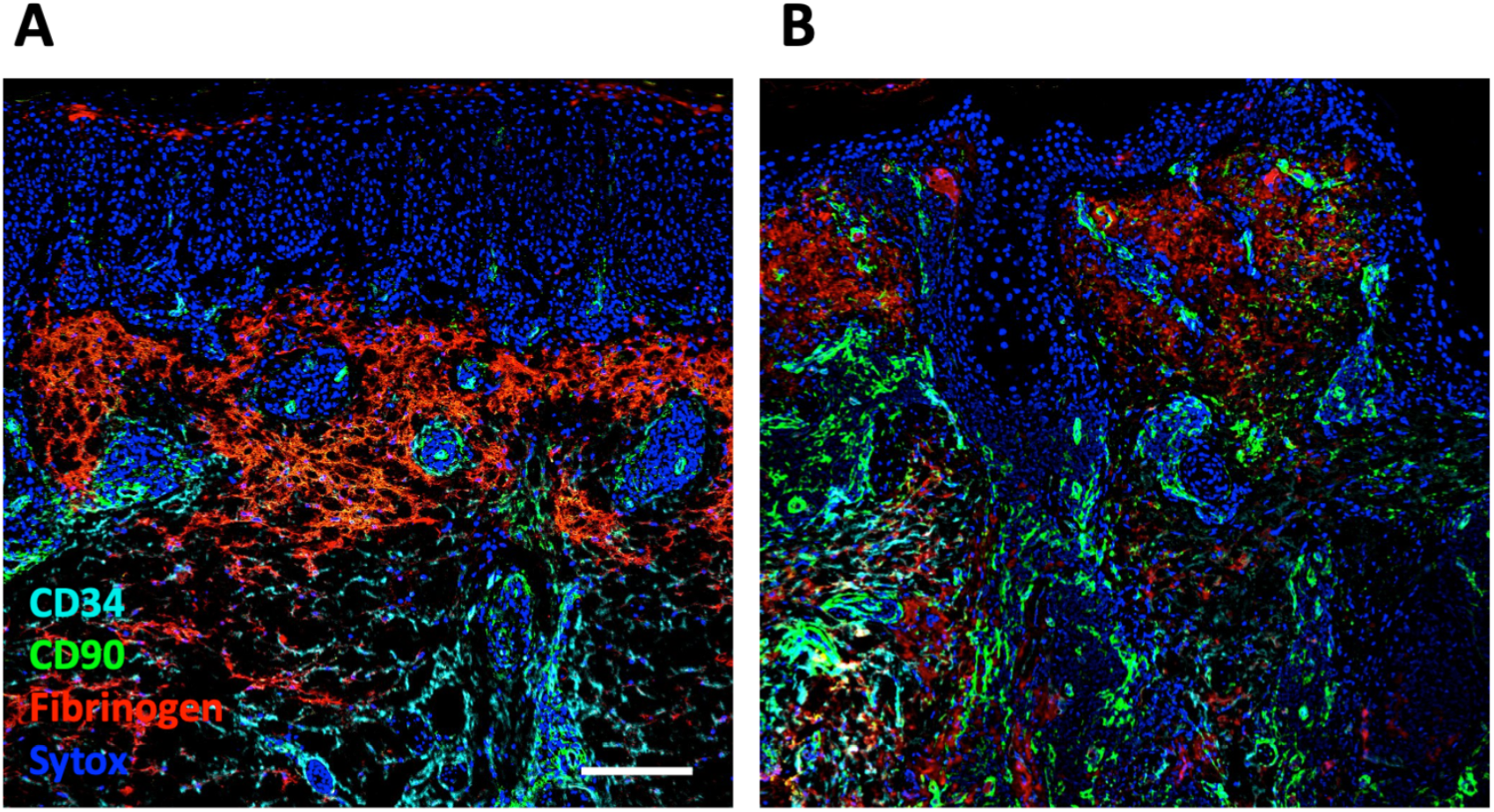
A). Comparison of fibrinogen deposition in inflamed skin of subacute (A) and chronic (B) lupus. Characteristic of cutaneous lupus, but atypical in SSc, is the epidermal hyperplasia and intensive perivascular leukocytic infiltration. Staining was for CD34, CD90 (pericytes and perivascular adventitial fibroblasts), Fibrinogen-γ and Sytox (nuclei). Scale bar is 200 µm.

Scant fibrin deposition was seen in about 20-25% of the dcSSc and lcSSc biopsies. If fibrinogen deposition is a critical driver of SSc, then should any stain-negative samples be present? Lack of staining by the pan-fibrinogen polyclonal antibody ruled out technical reasons. What distinguishes these fibrin-free skin samples is unclear, but they could represent patients with arrested disease. Nonetheless, fibrin deposition was extensive and widespread in both limited and diffuse forms in both early and late disease.

### Comparison to the inflamed skin in cutaneous lupus

Since fibrinogen leakage is a common feature of inflammation in general, we questioned whether the excessive deposition in SSc was unique or simply exemplary of inflamed tissue. Biopsies from cutaneous lupus patients were examined for comparison. In prior work, fibrin deposition had been reported at the dermo epidermal junction in cutaneous lupus and indeed we found fibrin in most cases (61). Interestingly, fibrin deposition was excluded from the infiltrated regions in chronic (discoid, N = 6) and subacute cutaneous (n = 4) lupus as well as lichen planus (n = 2) in all samples examined. In these tissues, the inflamed regions are characterized by a CD90+ immunofibroblast network and as expected from our prior work, this network in lichen planus as in lupus was also VCAM+ (not shown) (51). Similarly, the expanded CD90 positive perivascular adventitial compartment in SSc was largely free of fibrin (e.g., Figure 1B). Subacute and chronic spongiotic dermatitis biopsies also exhibited fibrin deposition (data not shown). The presence of fibrin in the deep reticular dermis in SSc and some cases of morphea differed from cutaneous lupus and lichen planus (e.g., Figure 1A,C, Figure 2B, and Supplemental Figure 5). When compared to cutaneous lupus, the cellular infiltration in SSc was minor. Thus, fibrin deposition in SSc appears to differ from that in the more inflamed lupus or lichen planus skin.

### Fibrin deposition is less evident in other pathological scarring events

Physiological and pathological skin scars e.g., keloids and hypertrophic scars were examined to determine whether fibrin deposition is a general characteristic of skin remodeling/scarring. The central fibrotic regions of both keloids and hypertrophic scars were fibrin free (n = 6 for each type, Supplemental Figure 6). Pathological scars are complex with central fibrotic regions often with more active regions around the edges. Fibrinogen was observed on the edges of some of these scars. Notably, the fibrin deposition in SSc skin was more extensive than that in pathological hypertrophic and keloid scars. Tumor vasculature can be leaky (27), yet biopsies from dermatofibroma (n = 2), dermatofibrosarcoma protuberans (n = 2), neurofibroma (n = 3) and atypical fibroxanthoma (n = 5) tumors did not show excessive fibrin deposition when compared to SSc (data not shown, note of the 5 atypical fibroxanthoma tumors examined, 1 was fibrin positive).

### Presence of the Fibrinogen-α_E_C domain differentiates the age of vascular leaks in physiological skin repair

Fibrin deposition occurs immediately following trauma from tissue excision, leading to hemostasis and wound containment. This fibrin-centric provisional matrix is essential to initiate wound repair, yet eventual dissolution of the clot via fibrinolysis is also required for proper healing (62). 1-3% of fibrinogen-α chain transcripts contain an alternative splice variant encoding a 34 kDa C-terminal extension called fibrinogen-α_E_C (Fib-α_E_C) and this domain is quite evolutionarily conserved (63). This Fib-α_E_C domain is readily cleaved from the parent fibrinogen yet the domain itself is relatively resistant to proteolysis. Curiously, the function of this domain remains unknown although a recent study showed that Fib-α_E_C was involved in clotting in zebrafish (64). The reported facile cleavage of the Fib-α_E_C domain from fibrin raised the possibility that its presence on the network could provide insight into the age of a fibrin deposit. We postulated that young clots would retain both Fib-α_E_C and Fib-γ, whereas older clots would lack Fib-α_E_C. To explore this hypothesis, we imaged Fib-γ, Fib-α_E_C and Factor XIIIA (F13A). F13A was chosen because it plays a critical role in fibrinolysis and inflammation via crosslinking fibrin and stabilizing the clot (65). F13A is also a macrophage marker that tracks with CD163 expression in repair-oriented macrophages and, indeed, tissue resident macrophages are considered the major source of the F13A present in the blood (66, 67). We had previously studied a series of reparative scars and here these samples were utilized to look at fibrin deposition at various time points post wounding (68). Colocalization of Fib-α_E_C and Fib-γ in the network was observed during the first week post excision indicating a roughly 1-2 week time frame for cleavage and separation of the Fib-α_E_C domain from the primary meshwork. The earliest sample at day 2 exhibited regions of dominant Fib-α_E_C that appeared as blood clots (as opposed to an extravascular erythrocyte/platelet free fibrin network). By 3-5 weeks, deposits shifted to being largely Fib-γ dominated or containing heterogenous regions presumably reflecting varying degrees of Fib-α_E_C release (Figure 5). Total clearance of Fib-γ was variable but required at least 5-14 weeks. Notably, we had observed in our previous work that a sample taken 2 days after excision contained regions with intact epithelium 3 mm distant from the excision, yet the dermis in this distant region had undergone the CD34-to-CD90 fibroblast transition (68). It was speculated that there was a rapid and extensive lateral propagation of damage (68). Here we found that fibrin deposition was also widespread throughout these same propagated regions raising the possibility that fibrin deposition could be connected to the fibroblast activation. This analysis included 11 reparative scar biopsies and hence represents a crude estimate of the time frame for clearance. Nonetheless, this wound analysis indicates that abundance of Fib-α_E_C in a deposit is inversely proportional to the age of the event.

**Figure 5:**
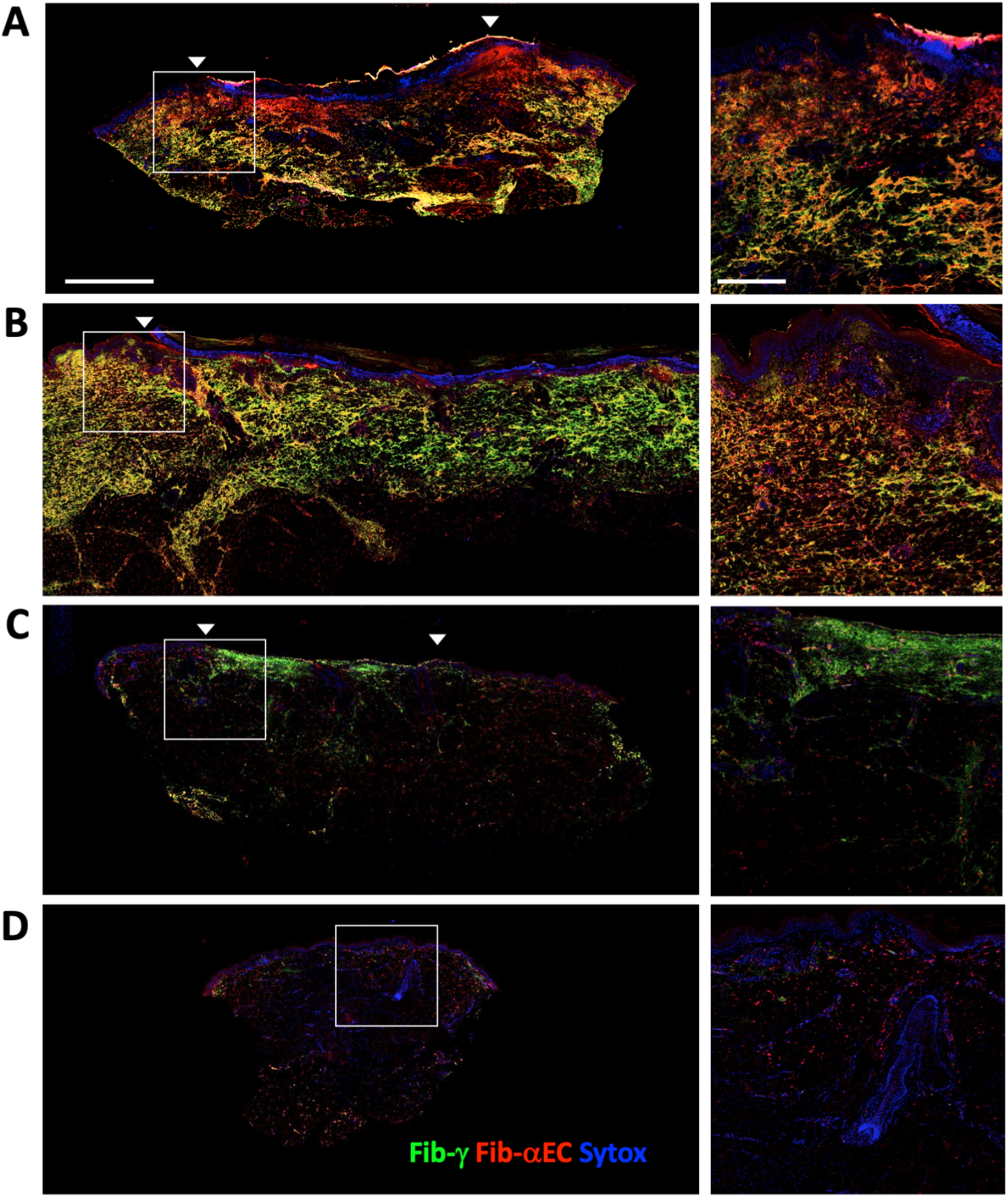
Relationship between bulk fibrin deposition (Fibrinogen-γ) and the presence of the Fib-α_E_C domain at (A) 2 days, (B) 7 days, (C) 98 days and (D) 151 days post excision wound. Arrows show sites of original excision and sytox identifies nuclei. Strong sytox nuclear staining at early time points is in an infiltrated region prior to re-epithelialization. Right panel shows a higher magnification with scale bars of 2 mm (left) and 500 µm (right).

### Fibrinogen-α_E_C domain reveals heterogeneity in fibrin deposition in SSc

We postulated that the appearance of Fib-α_E_C+ deposits would indicate active leaks in SSc with the expectation that long-standing disease would not show evidence of active leaks. We examined 12 healthy control, 4 lcSSc and 11 dcSSc biopsies and observed considerable inter-patient heterogeneity with a mixture of both Fib-γ single positive (or Fib-γ dominant) and Fib-γ+ Fib-α_E_C+ double positive regions in some tissues while other biopsies had solely the single positive Fib-γ pattern (Figure 6A-C). Examples of both patterns were seen in early and long-established disease. Curiously, Fib-α_E_C dominant fibrin often lined the lumen of the blood vessels (e.g., Figure 6B). While dcSSC disease generally plateaus or improves, a subset of these patients will continue to progress, thus, the presence of Fib-γ+ Fib-α_E_C+ deposits may be a novel indicator of active disease in SSc (58).

**Figure 6:**
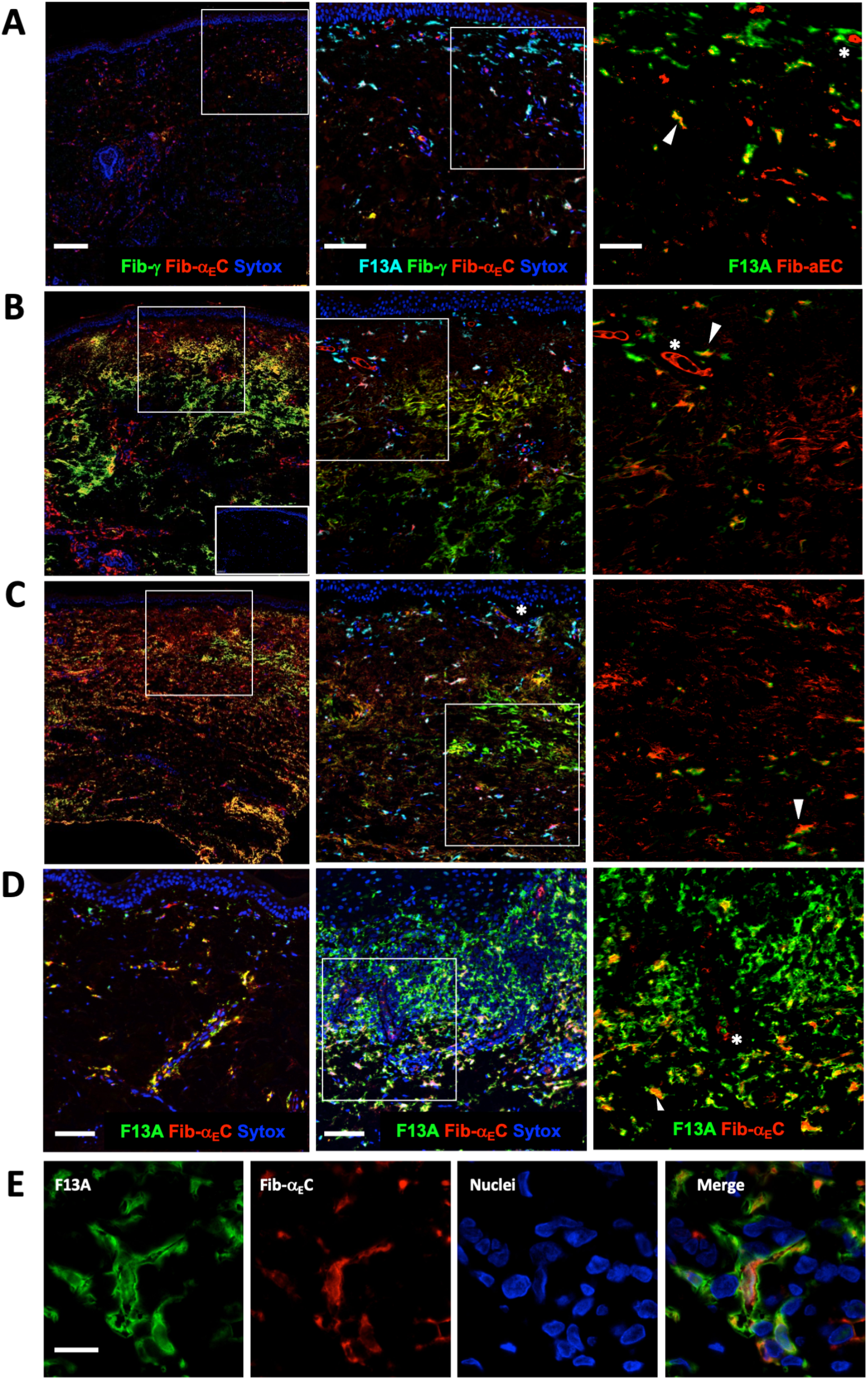
Relationship between bulk Fib-γ deposition and presence of the Fib-α_E_C domain in (A) healthy control, (B) early dcSSc (2 month duration, mRSS 42), (C) late dcSSc (+20 years duration, mRSS 33). Boxed regions are expanded on the right side and include the macrophage marker F13A. In the right most panels, the Fib-α_E_C signal was enhanced to emphasize colocalization with F13A. Insert in lower B is a secondary antibody alone control. (D) Colocalization of Fib-α_E_C domain with F13A+ macrophages in a healthy control (left) and lichen planus (middle) with an expanded lichen planus region on the far right. (E) High power image of a typical F13A+ macrophage with more central Fib-α_E_C domain largely surrounded by F13A. Asterisks mark Fib-α_E_C+ blood vessel lumen and arrows show examples of Fib-α_E_C/F13A double positive macrophages. Scale bars are 200 µm (left), 100 µm (mid) and 50 µm (right) (A-C), 100 µm (D) and 10 µm (E).

Surprisingly, some F13A+ macrophages in skin samples contained Fib-α_E_C (Figure 6A-D), yet consistently lacked detectable Fib-γ. Fib-α_E_C also co-stained CD163+ macrophages, which was expected since CD163 and F13A largely identify similar populations of dermal macrophages (data not shown) (69). F13A+ Fib-α_E_C+ macrophages were found in normal, SSc, lichen planus and DLE sections; with the notable paucity of Fib-α_E_C+ macrophages in the highly leukocyte infiltrated regions of lichen planus (Figure 6D) and cutaneous lupus (not shown). We hypothesize that upon Fib-α_E_C release, this domain accumulates in macrophages perhaps via Fib-α_E_C binding to integrin α_V_β_3_ (63). Abundant Fib-α_E_C negative macrophages in the infiltrated regions are likely cells that had recently migrated into the perivascular compartment from the blood and have not yet ingested Fib-α_E_C. F13A+ dermal macrophages are known to be long lived (70) and our observations are consistent with a model where F13A+ resident macrophages accumulate Fib-α_E_C following vascular leaks. Notably, neonatal foreskin only contained Fib-α_E_C-F13a+ macrophages consistent with an insufficient time or prior leakage for Fib-α_E_C accumulation (data not shown). Therefore, Fib-α_E_C+ accumulation may be an indicator of extended macrophage residence.

### Fibrin deposition in relationship to other events in SSc

We questioned whether the appearance of fibrinogen deposition correlated with other known events notably the loss of CD34 protein expression by dermal fibroblasts in SSc (48, 50). More recently, loss of CD34 and gain of smooth muscle actin (SMA) were identified as robust markers of disease severity (49). Using morphometric analysis, we found the pattern of regional CD34 and fibrinogen expression in the PRESS biopsies was consistent with an inverse relationship between high fibrin levels and low fibroblast CD34 levels (Supplemental Figure 7). Plotting the regional density of fibrin vs CD34 appears to be a vivid indicator of SSc disease.

Next, we probed bulk RNA sequencing data available from paired biopsies from the PRESS collection (60). We had previously reported that the ratio of bulk CD90 to CD34 RNA tracked with the extent of skin disease based on MRSS (48). The CD90/CD34 ratios from both the early (PRESS) and the (GENISOS) disease datasets distinguish SSc patients from controls and importantly the increase occurs early in the disease course (PRESS Cohort) (Supplemental Figure 8A). The CD90/CD34 ratio increase in these two new cohorts tracks roughly with MRSS scores but not obviously with disease duration (Supplemental Figure 8B,C). The change in this ratio is largely accounted for by increased CD90 RNA (Supplemental Figure 8D-F) and is likely driven by both fibroblast transitioning to a CD90 positive state as well as an expanded perivascular adventitial compartment containing CD90 positive fibroblasts (51, 71). The exact relationship between protein expression as detected by immunohistology of the highly glycosylated CD90 and CD34 molecules and their RNA levels remains incompletely defined (48, 51, 68, 72, 73).

To further interrogate the PRESS samples, a heat map was created using histology scores, cell type RNA signature scores as defined previously, and the CD90/CD34 RNA ratios (Figure 7) (60). Forced ranking based on the CD90/CD34 RNA ratio revealed several points. First, all disease and control subjects were correctly grouped and the CD90/CD34 ratio correlated well with both local and global mRSS scores and a myofibroblast histology score (Spearman r = 0.73 and 0.79). Given the documented increases in expression of the fibrinolysis inhibitor PAI-1 in SSc, we examined RNA expression levels of both SERPINE1 and 2, i.e., PAI-1 and PN-1, etc. Expression of SERPINE2 was more abundant than SERPINE1 and both genes correlated with CD90/CD34 ratio changes. Second, fibrin deposition appears in the PRESS SSc cohort and hence is an early event. Third, fibrin presence did not correlate with the CD90/CD34 RNA ratio. Fourth, fibrin deposition correlated best with the CD4 and DC signatures suggesting a relationship between the local immunology and early vascular leakage. Overall, the CD90/CD34 ratio changes correlated well with the fibrotic manifestations.

**Figure 7:**
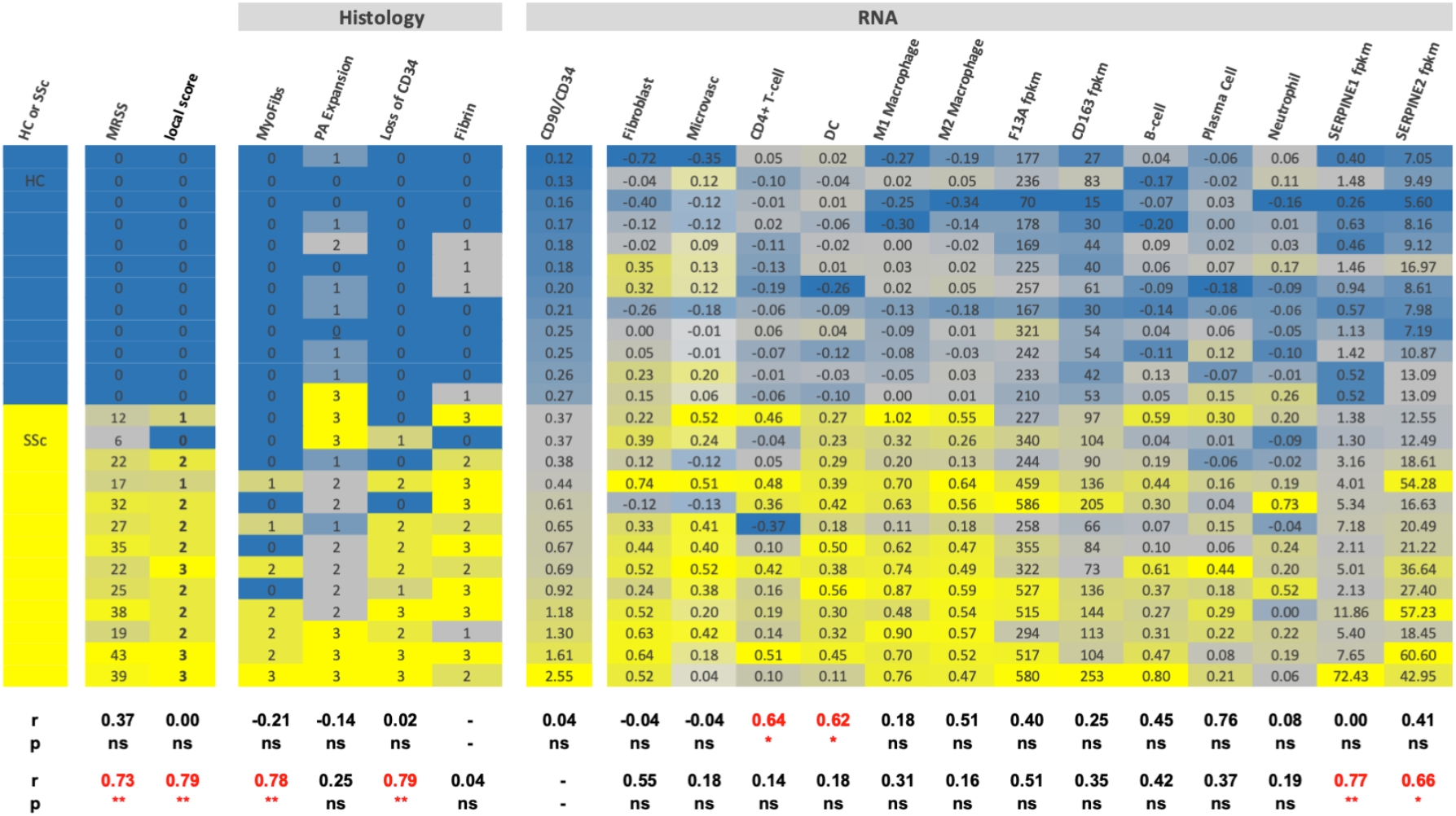
A). Heat map illustrating the correlations between histological fibrinogen deposition and various RNA cell signatures or individual gene expression in the PRESS cohort patients based on bulk RNA seq data. PA expansion refers to increased CD90+ perivascular adventitial compartment. Samples were ranked according to the bulk RNA CD34/CD90 ratio. Spearman correlation coefficients were determined relative to the fibrin histology score (upper row) and the CD90/CD34 RNA ratio (lower row) and only SSc patient data was used in the correlation analysis. The quantal histology scoring was treated as a continuous variable for this analysis.

Macrophage infiltrates are an early feature of SSc skin disease (74, 75). Macrophages have a complex relationship with wound healing, with both positive and negative effects possibly related to whether they are inflammatory blood-derived or tissue resident macrophages (76). We reexamined whether macrophages either globally or specifically co-localized with fibrin deposition in the PRESS samples. Visually, F13A+ macrophages did not show an obvious spatial co-localization with regions of fibrin deposition with the caveat that in many samples, fibrin deposition was too widespread (Figure 6A-D). Correlation between CD90/CD34 RNA ratios and bulk macrophage RNA signatures did reach significance. Overall, these data indicate that both CD90/CD34 RNA ratio changes and vascular leakage i.e., fibrin deposition are relatively early events.

## Discussion

Here we show that dermal extravascular fibrin deposition is extensive in SSc, a result that would be difficult to observe in a transcriptomic analysis as fibrinogen genes are expressed in the liver. Likewise, the presence of intravascular and external blood clotting due to the biopsy excision procedure complicates bulk proteomic approaches. The extensive fibrin accumulation occurs early in the disease course and is retained in advanced disease. This cutaneous observation is consistent with prior blood analyses indicating general hypercoagulation and decreased fibrinolysis in SSc (44). We speculate that early unknown triggers lead to vascular leakage and extracellular fibrin accumulation which in turn drives downstream chronic tissue remodeling.

Given the known symptom of “puffy” hands and fingers in early SSc, the observation of extensive fibrin deposition could provide a molecular link between early events and the eventual fibrotic manifestations. Puffy hands were recently highlighted as an important criterion of very early disease (77). As vascular leak and edema is one of the cardinal signs of inflammation, a central question is whether the extensive fibrin deposition observed here in SSc is atypical. Several observations suggest the extensive fibrin deposition is indeed unusual. First, dcSSc disease has an early inflammatory phase during the first 1-3 years followed by slowed progression or even improvement (57–59). Hence, the continued presence of fibrin deposits in the late phase is not tracking simply with the inflammatory phase. Second, fibrin deposition in SSc is relatively widespread throughout the dermis including in peri-eccrine and adipose regions, whereas in the inflamed tissue in cutaneous lupus and lichen planus, deposition is largely in the papillary dermis or epidermal-dermal interface regions. Third, the relative lack of fibrin deposition in pathological scars and cutaneous mesenchymal fibrous tumors differentiates SSc. While SSc is considered an inflammatory disease, the relative paucity of F13A+ macrophages in SSc when compared to the heavily infiltrated regions in lichen planus or cutaneous lupus raises the possibility that the level of inflammation in SSc may be inadequate to drive resolution of the fibrin deposits.

Given the persistent nature of fibrin deposition in SSc, a second question is whether deposition is continuous, or its removal is impaired. It was in this light that we sought to utilize the presence the Fib-α_E_C domain on the fibrin deposits as a marker of recent leakage. Analysis of a series of timed reparative scar biopsies was consistent with a relatively fast loss of network-associated Fib-α_E_C and a considerably slower loss of the Fib-γ reactive epitope. We found Fib-α_E_C rich deposits variably in both early and late SSc skin, albeit with local heterogeneity. Thus, use of this tool suggests that there may be ongoing vascular leak even in some late-stage patients, perhaps indicative of active disease and/or continued progression. Fibrin with a tangle-like appearance in mid to deep dermal regions in SSc lacked the Fib-α_E_C domain and hence appear to be older consistent with impaired removal. It is possible that local heterogeneity in the proteolytic milieu could impact Fib-α_E_C release rather than fibrin network longevity. Nonetheless, the determination of the status of disease progression in SSc has been especially challenging and this new Fib-α_E_C tool may provide novel insight. Knowledge of the rates of clearance of fibrin networks in human tissue wounds, tumors or inflammatory disease is limited and hence this method could find application in many settings outside of SSc.

The persistence of the Fib-α_E_C domain in resident-like macrophages is an intriguing observation that raises many questions. Relatively few proteins remain resistant to digestion in macrophages or even in interstitial spaces with examples including amyloid forming proteins, tau protein, etc. The recently reported retention of α-synuclein within microglia without breakdown may be an instructive example (78). It was speculated that following proteolytic release by matrix metalloproteinases or plasmin, Fib-α_E_C may signal between fibrin deposition and the inflammation/tissue repair systems (63). Its presence in normal tissue indicates that macrophage Fib-α_E_C retention is a normal physiological process perhaps reflecting accumulation following routine dermal micro-trauma. The lack of this domain in neonatal foreskin macrophages supports this model. Curiously, bulk fibrin deposition was not observed in the heavily macrophage infiltrated perivascular adventitial compartment in both the inflamed settings of lichen planus and lupus as well as in the expanded perivascular adventitial compartments typically observed in SSc. As heavily macrophage infiltrated perivascular regions in lichen planus and lupus tended to lack the Fib-α_E_C domain, it is likely that these perivascular macrophages are new immigrants from the blood that have not had time to accumulate Fib-α_E_C. Perhaps these perivascular regions contain excess fibrinolytic machinery or, alternatively, pre-formed fibrin deposits were pushed out upon leukocytic infiltration. Whether Fib-α_E_C is simply a long-lived fibrin breakdown product, an active component of proper tissue repair or even a contributor to the pathology in certain disease settings warrants further investigation.

A third major question is whether the chronic fibrin deposition contributes directly to the downstream fibrosis in SSc. Clot formation and its removal via fibrinolysis are widely recognized as a central biological process as well as a prerequisite for successful tissue repair following injury (11, 12, 62, 79). In rodent models and in human pleural injury, a lack of fibrin clearance hinders wound repair culminating in fibrosis i.e., consistent with a linkage between chronic vascular leak and pulmonary fibrosis (12, 80, 81). Fibrinogen is a multi-faceted protein, capable of engaging the innate immune machinery and the vasculature as well as binding growth factors and latent TGF-β (23). Studies in multiple systems support causality in inflammatory and neurological diseases (23, 79, 82). Mice genetically deficient in fibrinogen or genetic inhibition of its interaction with αMβ2 with impaired inflammatory activity are protected in a wide range of disease models including rheumatoid arthritis, periodontitis, multiple sclerosis and Alzheimer’s disease, suggesting that fibrinogen is a key driver of inflammation and fibrosis in tissues (15, 23, 83–85). Other components of the coagulation cascade also participate in disease pathogenesis through fibrinogen-dependent and independent mechanisms (19, 21, 79, 86, 87). F13A itself plays a role in fibrinolysis and genetic mutations in this gene drive familial forms of dermatofibroma (88). Another intriguing possibility lies in fibrin’s role in forming the “provisional matrix” for wound repair. This matrix could be a form of altered “stiffness” resulting in pro-fibrotic mechano-sensing and fibroblast reprogramming (89, 90). It is perhaps relevant that engineered fibrin hydrogels have been extensively investigated in tissue repair settings e.g., burn repair, post-surgical healing and orthopedics.

The role of fibrin deposition in murine lung fibrosis has been studied using different genetic backgrounds with complex results. Intratracheal bleomycin induces direct epithelial damage with intra-alveolar fibrin accumulation (91). Bleomycin-induced lung fibrosis still occurred in the genetic absence of fibrinogen, albeit aspects of the pathology were altered (92, 93). In contrast, removal of fibrinolysis components, i.e., loss of tissue plasminogen activator or gain of PAI-1 both exacerbated fibrosis (94). Activation of PAR1 by thrombin is important in a modified bleomycin lung fibrosis model where bleomycin induced fibrosis is exacerbated by enhanced vascular leakage (16, 95). In the skin, fibrosis can be directly triggered by injection of fibrin into a tissue plasminogen activator deficient mouse, i.e., a setting of impaired fibrinolysis (20). It is possible that the role of fibrin in the skin, liver, kidney and lung in animal models of fibrosis depends on tissue-specific pro-coagulant and fibrinolytic mechanisms in different anatomical compartments with different cellular compositions e.g., airways and epithelium vs interstitial spaces.

We attempted to determine whether fibrin deposition correlated with the dramatic and well-documented loss of fibroblast CD34 staining in SSc (48–50). Morphometric analysis of the PRESS cohort samples was suggestive of an inverse relationship wherein regional fibrin appearance roughly paralleled CD34 loss. Furthermore, in an early reparative wound, widespread fibrin deposition at considerable distance from the primary excision was accompanied by the loss of CD34 expression. Fibroblasts within pathological scars have undergone the CD34 transition, and there, the paucity of fibrin deposition would argue against fibrin involvement in scar formation. However, the active remodeling regions may lie at the tissue interface or fibrinolysis may be more functional in these scars. We strove to find parallels between skin bulk RNA profiles and the fibrin deposition. The correlation between the CD90/CD34 RNA ratio and the mRSS clinical score was confirmed with two new datasets and importantly this change occurs early in the disease course (48). In the PRESS cohort, this ratio tracked with myofibroblast appearance consistent with another recent analysis (48, 49). While the elevation of SERPINE1 (PAI-1) has received considerable attention in SSc, SERPINE2 (protease nexin-1, PAI-2) was also elevated and both genes tracked with the CD90/CD34 RNA ratio. SERPINE2 can limit fibrinolysis and is potentially involved in cardiac fibrosis (96). Despite the apparent regional association between fibrin deposition and loss of CD34 display, fibrin presence did not track with the CD90/CD34 RNA ratio. It is possible that local regional interactions are not robustly manifested in the bulk RNA data or the connection between these two components is not direct. Importantly, histological scoring of fibrin deposition correlated with increased CD4 T cell and dendritic cell RNA signatures suggesting a relationship between T-cell based immunology and early vascular leakage. This observation is consistent with the role of fibrinogen in promoting autoimmunity and antigen presentation in the brain (97). In SSc, it is unclear whether this correlation reflects a primary immunological induction of vascular leakage or is a consequence of fibrin deposition (97). These analyses are consistent with a chronology where early T-cell/DC involvement occurs during vascular leak and fibrin deposition. A decrease in fibroblast CD34 expression, myofibroblast appearance and tissue fibrosis as reflected by mRSS scoring lie downstream. The known proinflammatory role of fibrin in activation of innate immune cells could provide linkage between these early and later sets of observations (97).

The limitations of this study include variables associated with tissue collection, indeterminant lesioned status especially in lcSSc biopsies and the limited number of paired PRESS biopsies for histology and RNA analyses. We cannot exclude the possibility that the pro-fibrotic myofibroblast phase can occur early, is transient and was already resolved in some early disease cases affecting our assessment of the order of events. Interpretation of the data is limited by a dearth of knowledge surrounding the longevity of fibrin deposits in normal wound healing or other injury settings as well as robust clinical indicators of active vs quiescent SSc.

In summary, we have shown here that extensive and persistent fibrin deposition occurs in SSc and have outlined a novel approach to explore the relative age of a fibrin deposit. The possibility that long-term fibrin presence in the skin serves as a chronic bridge to tissue remodeling has ramifications for a better assessment of SSc disease heterogeneity, staging and progression as well as the development of novel therapeutic intervention strategies. A first-in-class fibrin immunotherapy neutralizing fibrin-induced immune activation without adverse effects in hemostasis protects from neurologic disease in mouse models of multiple sclerosis and Alzheimer’s disease (98).

## Methods

### Patient Skin Biopsies

Skin biopsy specimens were obtained from the Arthritis and Autoimmune Diseases Center at Boston University Medical Center as described previously (99). All patients with diffuse SSc (dcSSc) or limited SSc (lcSSc) met the American College of Rheumatology criteria for SSc (100). The study was conducted under a protocol approved by the Institutional Review Board of Boston University Medical Center, and written informed consent was obtained from all subjects. Skin punch biopsy specimens were obtained from lesioned regions of the dorsal mid forearm of SSc patients. Healthy control subjects had no history of skin disease. Sections were also accessed from an additional PRESS cohort of skin biopsies from early dcSSc patients and healthy control subjects along with adjacent biopsies for bulk RNA sequencing as previously described (60). Additional bulk RNA sequence data was obtained from the entire PRESS and GENISOS cohorts (60, 101). Patient characteristics are defined in Supplemental Table 1. For comparative purposes, further skin sections were provided by the Skin Pathology Laboratory (Dermatopathology Section Boston University) and classified as representative of the disease by the pathologists Drs. J. Bhawan and J. Ho. These biopsies included (n examined): morphea (8), subacute (4) and chronic (discoid) (6) lupus erythematosus, lichen planus (2), dermatofibroma (2), dermatofibrosarcoma protuberans (2), neurofibroma (3) and atypical fibroxanthoma (5) as well as skin scars including normal reparative (11), hypertrophic (6) and keloid (6) which have been analyzed in this lab in previous studies (48, 51, 68).

**Table 1:** Fibrinogen Scores at Various Disease Stages.

|  | Duration<br>(Years) | n | Mean | SD | P to<br>control |
| --- | --- | --- | --- | --- | --- |
| **All Data** |  |  |  |  |  |
| Controls | - | 28 | 0.50 | 0.64 | - |
| DsSSc |  |  |  |  |  |
| Total | - | 69 | 1.73 | 1.07 | <0.0001 |
| Early | 0-2 | 43 | 1.77 | 1.11 | <0.0001 |
| Mid | 2-6 | 14 | 1.64 | 0.93 | 0.012 |
| Late | > 6 | 12 | 1.67 | 1.15 | 0.016 |
| LsSSc |  |  |  |  |  |
| Total | - | 21 | 1.52 | 1.12 | 0.009 |
| Early | 0-4 | 11 | 1.56 | 1.04 | 0.059 |
| Late | >4 | 10 | 1.5 | 1.27 | 0.096 |
| Morphia | - | 8 | 1.50 | 0.53 | 0.220 |
| **PRESS Cohort** |  |  |  |  |  |
| Controls |  | 12 | 0.33 | 0.49 | - |
| DcSSc |  |  |  |  |  |
| Early |  | 13 | 2.08 | 1.04 | 0.0001 |
| **Percentage of Patients with a Fibrinogen Score** |  |  |  |  |  |
|  |  |  | **Fibrinogen Score (%)** |  |  |
| **All Data** |  | **0** | **1** | **2** | **3** |
| Controls | - | 57 | 36 | 7 | 0 |
| DsSSc |  |  |  |  |  |
| Early | 0-2 | 22 | 12 | 37 | 30 |
| Mid | 2-6 | 14 | 21 | 50 | 14 |
| Late | > 6 | 17 | 33 | 17 | 33 |
| LsSSc |  |  |  |  |  |
| Early | 0-4 | 18 | 27 | 36 | 18 |
| Late | >4 | 30 | 20 | 20 | 30 |

### Immunohistochemistry

Sections from formalin-fixed paraffin embedded skin were processed as previously described by deparaffinization and antigen heat retrieval in Tris-EDTA buffer pH 9 (51). Endogenous peroxidase and phosphatases were blocked sequentially with 3% H2O2 and then BLOXALL (Vector Laboratories) followed by blocking with 10 mg/ml BSA. Antibodies used in this study are described in Supplemental Table 2. Two color conventional staining was performed using mouse anti-fibrinogen-γ combined with rabbit anti-CD34 and developed first with ImmPress anti-mouse HRP polymer and AMEC detection followed by anti-rabbit ImmPress AP polymer and HighDef® blue AP stain (Enzo) effectively as described (48). In some cases of negative fibrinogen staining with the mouse anti-fibrinogen-γ antibody, we questioned whether the epitope for the Fib-γ mAb in the N-terminal region of the protein was variably present due to altered sensitivity to antigen retrieval, to the extent of fibrin cross-linking or the extent of the degradative fibrinolytic process. Direct comparison between the monoclonal and the polyclonal anti-fibrinogen antibodies using serial sections did not reveal a major discrepancy indicating that some samples indeed had minimal fibrin deposition.

Four-plex immunofluorescence was carried out effectively as described using three tyramide based staining layers followed by quenching between each layer of bound HRP-polymer with 3% H2O2 (51). Dyes included Biotium CF®405S, CF®488A and CF®594 followed by mounting in fluid containing 10 nM of the nuclear stain Sytox (InVitrogen). Slides were imaged using an Olympus FLuoView confocal microscope using the Alexa 405, 448, 594 and 647 (for Sytox) channels. Bleed through between the 594 and 647 channels was minimal with these optics. Some slides were also scanned using the Vectra Polaris Imaging system followed by spectral unmixing using the Akoya Biosciences Inform software. Tiles were stitched with Indica Halo software. Morphometric analysis of fibrinogen staining was carried out on the IHC PRESS images in ImageJ as described previously (51) with color deconvolution (fast blue/fast red settings) on a user-defined region of interest (ROI) that excluded the epidermis, hair follicles, sebaceous and eccrine glands, tissue rips and biopsy edges followed by standardized thresholding and pixel counting. Vessels with intravascular fibrin clots were included. Average area in the ROI was 55 +/- 15% of the total image field (2.4 mm^2^, 40x magnification) and the fibrin density was the percent fibrin positive pixels in the ROI. Co-localization analysis was performed with the ImageJ coloc2 program with Costes thresholding. Spatial co-localization of CD34 and fibrinogen was carried out using the 2-color IHC images obtained with the Mantra 2.0 microscope (Akoya Biosciences) with spectral unmixing using Inform software. With Halo software, a user defined ROI was overlaid to exclude margins and epidermis, constant thresholds were applied, and the ROI was overlaid with a 90 × 90 µm grid. Numbers of CD34+ or fibrinogen+ pixels were determined for each tile. Myofibroblast scoring of the PRESS biopsies was described previously and similar scoring for perivascular adventitial expansion has been outlined (51, 60).

### RNA Analyses

RNA correlations relied upon published bulk RNA-seq data from the PRESS and GENISOS cohorts as well as microarray data from the BU SSc patients and are deposited in the NCBI’s Gene Expression Omnibus database GSE130955 or as described previously (51, 60).

### Computational Analyses

All graphic and statistical analyses were carried out with Graph Pad Prism 9. Further details are included in each figure legend. Heat map was generated in Excel by forced ranking of the CD90/CD34 RNA ratio and color gradients employed conditional formatting (3-color, same parameters) for each column. Quantal histology scores were treated as continuous variables.

## Author Contributions

Histology analyses were conceptualized and carried out by JLB and RNA data analyses were performed by BS. JB and JH validated diagnoses and provided tissues from all non-SSc patients. AT and NC provided guidance for spectral-unmixing microscopy, slide scanning and downstream analysis. AMB gave useful guidance on SSc medical aspects as well as maintaining the current BU tissue repository. KA provided pivotal guidance on fibrin histology and interpreted data. SA coordinated sample collection from the PRESS and GENISOS cohorts. The study was funded by grants DoD W81XWH-61-1-0296 and the National Scleroderma Foundation (SA) and general funds from the BU Department of Microbiology. KA is supported by grants from NIH/NINDS R35 NS097976 and NINDS/NIA RF1 AG064926. BS is supported by the National Institute of Arthritis And Musculoskeletal and Skin Diseases (NIAMS) of the National Institutes of Health (NIH) under Award Number [K08AR081402] and by an Investigator Award from the Rheumatology Research Foundation (RRF).

## Acknowledgements

We thank the patients and investigators in the multicenter national PRESS registry (77), and the BU patients as well as Robert Lafyatis and Robert Sims for their contributions to the formation of the original BU SSc tissue repository. We are grateful to Aoife O’Connell and Hans Peter Gertje for microscopy and imaging help, Keith Wharton for critical advice and especially to Jen Gommerman who pointed out the use of fibrinogen to monitor vascular leakage in multiple sclerosis.

## Supplemental Data

**Supplemental Table 1:** Patient characteristics

**Supplemental Table 2:** Antibodies used in this study

**Supplemental Figure 1:** Comparison of Dako rabbit anti-Pan-fibrinogen and mouse anti-fibrinogen-γ mAb. Equivalence of two different antibodies to fibrinogen. Serial sections of a dcSSc biopsy (early disease), were stained with the fibrinogen-g mAb or the Dako rabbit anti-pan-fibrinogen polyclonal. Adipose/eccrine (middle) and fibrotic (left) regions are shown at higher magnification and fibrinogen leakage from small adipose and peri-eccrine capillaries is readily visible. Sections were counterstained for CD34 and the rabbit anti-CD34 mAb and the mouse anti-CD34 mAb exhibit some slight staining differences. Scale bar is 200 mm.

**Supplemental Figure 2:** Examples of the 0-3 scoring system for grading fibrin deposition. These 4 biopsies illustrate the general scoring system. The section with “0” has edge fibrinogen that results from bleeding during the core biopsy.

**Supplemental Figure 3:** Images of all the control biopsies analyzed from the PRESS cohort. Staining is fibrinogen-γ (brown) and CD34 (blue). Text insert includes the age, mRSS score, local MRSS score (0-3), the bulk RNA CD90/CD34 ratio and the fibrin density expressed as %positive pixels. Scale bar is 200 µm.

**Supplemental Figure 4:** Images of all the early dcSSc biopsies analyzed from the PRESS cohort. Staining is fibrinogen-γ (brown) and CD34 (blue). Text insert includes the age, MRSS score, local MRSS score (0-3), the bulk RNA CD90/CD34 ratio and the fibrin density expressed as %positive pixels. Scale bar is 200 µm.

**Supplemental Figure 5:** Examples illustrating the unique presence of fibrin in the deep dermis in SSc and morphea, but not in subacute or chronic cutaneous lupus. Fibrinogen levels in morphea are typically less than those in SSc.

**Supplemental Figure Fig 6:** Fibrin deposition in two examples of hypertrophic and keloid scars. In general, the central fibrotic regions including nodules lack fibrinogen deposition. When fibrinogen deposition is visible, it is on the outer edges of the scar region. There is mild inflammation in the fibrinogen rich region in this keloid scar.

**Supplemental Figure 7:** Analysis of Regional CD34 and Fibrinogen Expression. (A) IHC images were subjected to spectral unmixing, ROI definition, color thresholding, grid overlay (85 mm2) and the percent red or green pixels determined. These values are plotted in (B) where three examples of control and SSc (PRESS cohort) skin illustrate the range of results. In general, increased regional fibrinogen was accompanied by reduced CD34 levels.

**Supplemental Figure 8:** Relationship between CD34/CD90 RNA Ratio and Disease (A) Comparison of CD90/CD34 RNA ratio in the GENISOS (n = 33 control and 55 SSc) and PRESS (n = 33 control and 48 SSc) healthy control and dcSSc cohorts. Mann-Whitney U test. (B) Cross-sectional data showing the lack of a relationship between the ratio and disease duration. (C) Ratio correlates with increased MRSS scores, dotted line shows the mean of the control samples. (D) Receiver operating characteristic plots of all the PRESS and GENISOS CD90/CD34 ratio data (n’s as per A) which is largely driven by the CD90 RNA levels (E and F).

