## Supplemental materials for "Extensive and Persistent Extravascular Dermal Fibrin Deposition Characterizes Systemic Sclerosis"

### Slide 1
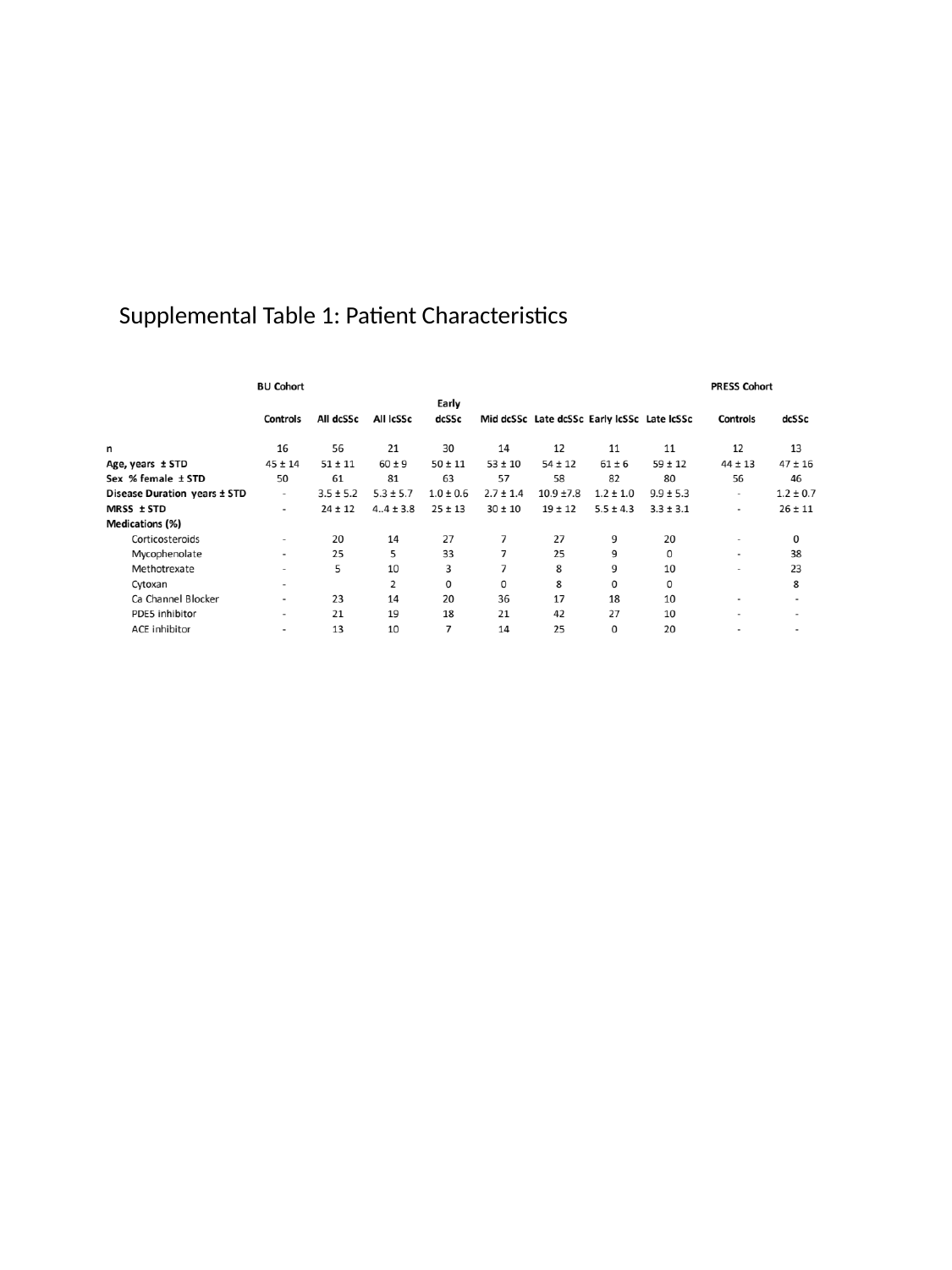

Supplemental Table 1: Patient Characteristics

### Slide 2
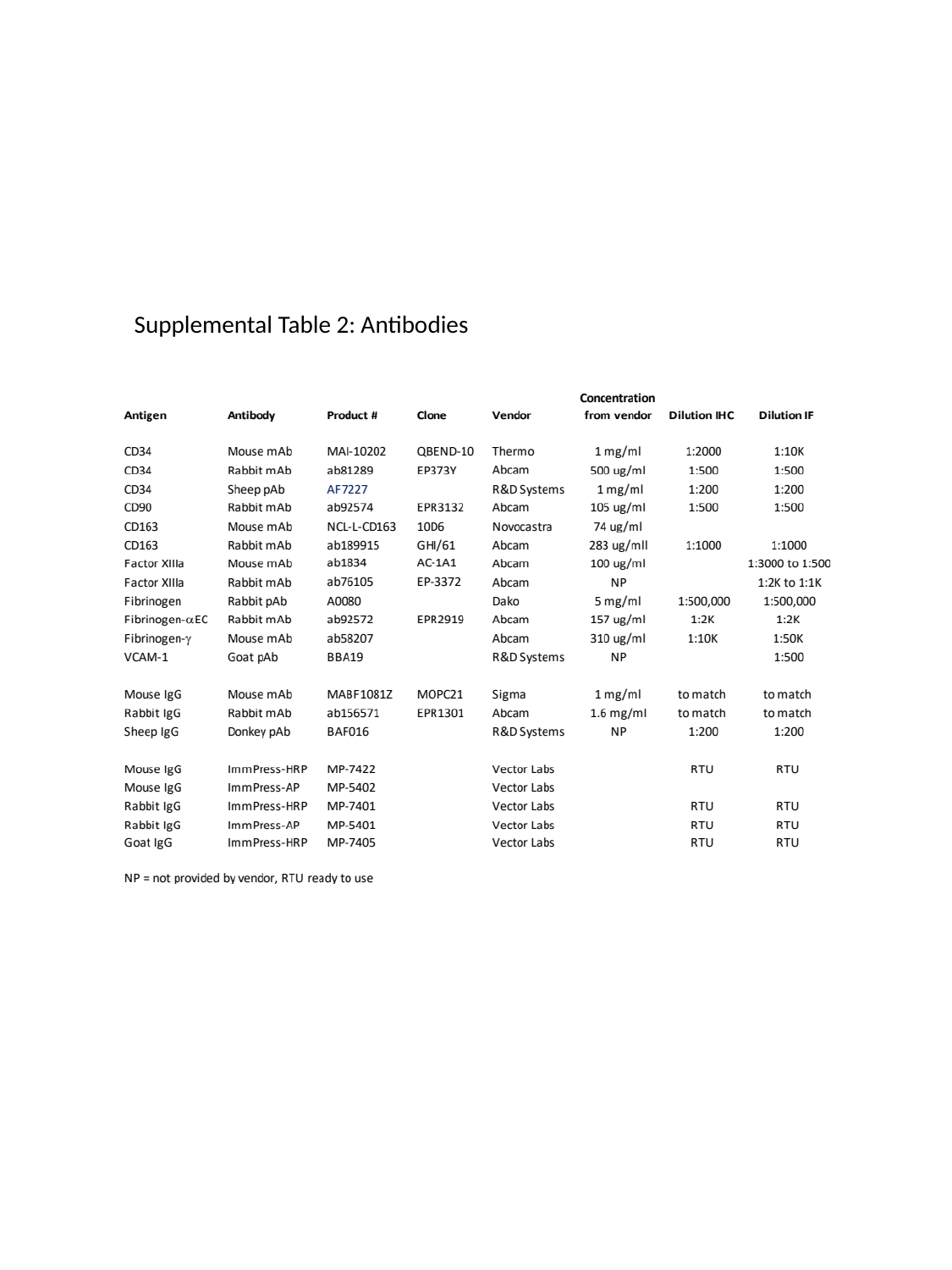

Supplemental Table 2: Antibodies

### Slide 3
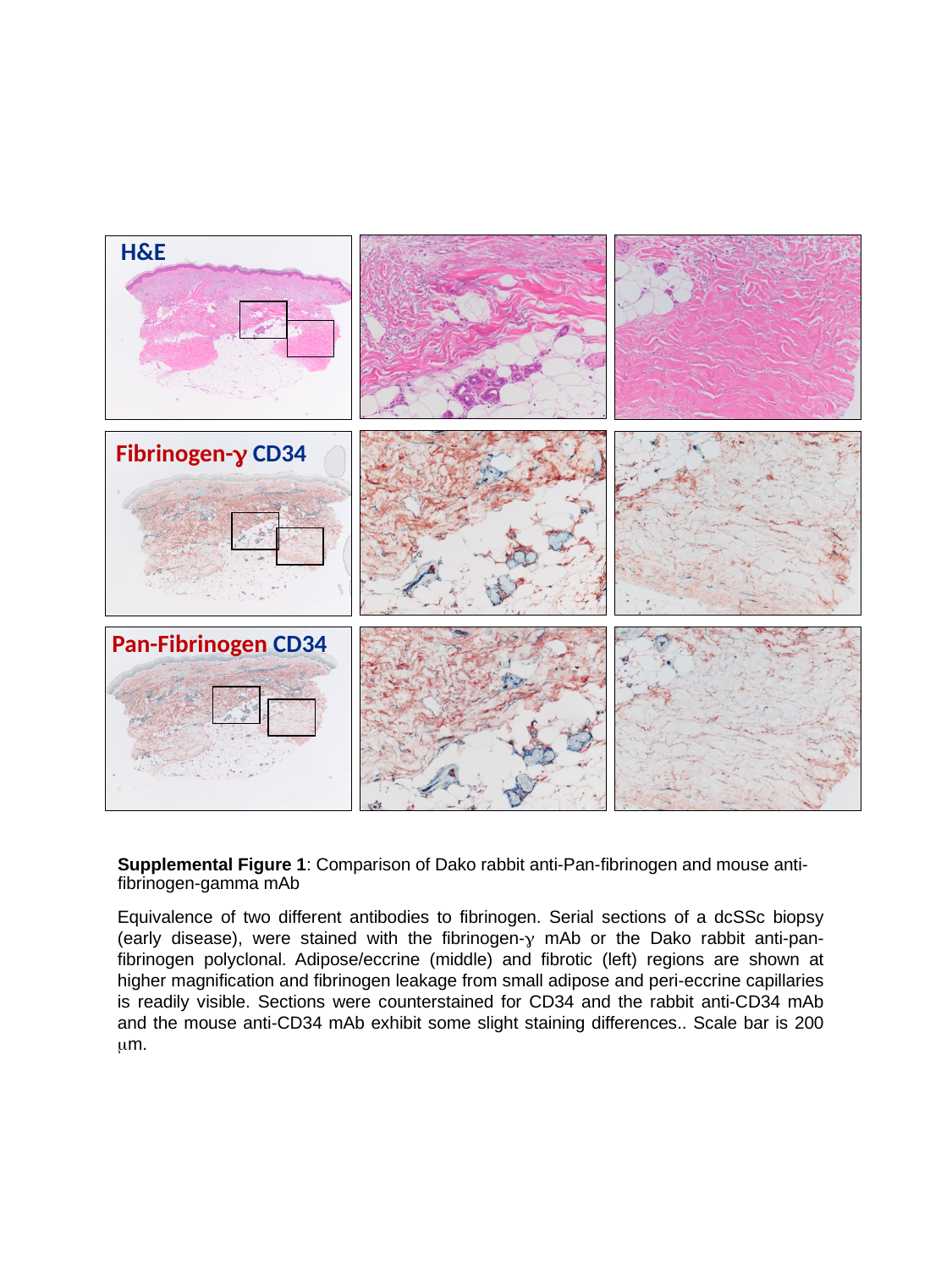

H&E
Fibrinogen-γ CD34
Pan-Fibrinogen CD34
#### Supplemental Figure 1: Comparison of Dako rabbit anti-Pan-fibrinogen and mouse anti-fibrinogen-gamma mAb
Equivalence of two different antibodies to fibrinogen. Serial sections of a dcSSc biopsy (early disease), were stained with the fibrinogen-γ mAb or the Dako rabbit anti-pan-fibrinogen polyclonal. Adipose/eccrine (middle) and fibrotic (left) regions are shown at higher magnification and fibrinogen leakage from small adipose and peri-eccrine capillaries is readily visible. Sections were counterstained for CD34 and the rabbit anti-CD34 mAb and the mouse anti-CD34 mAb exhibit some slight staining differences.. Scale bar is 200 μm.

### Slide 4
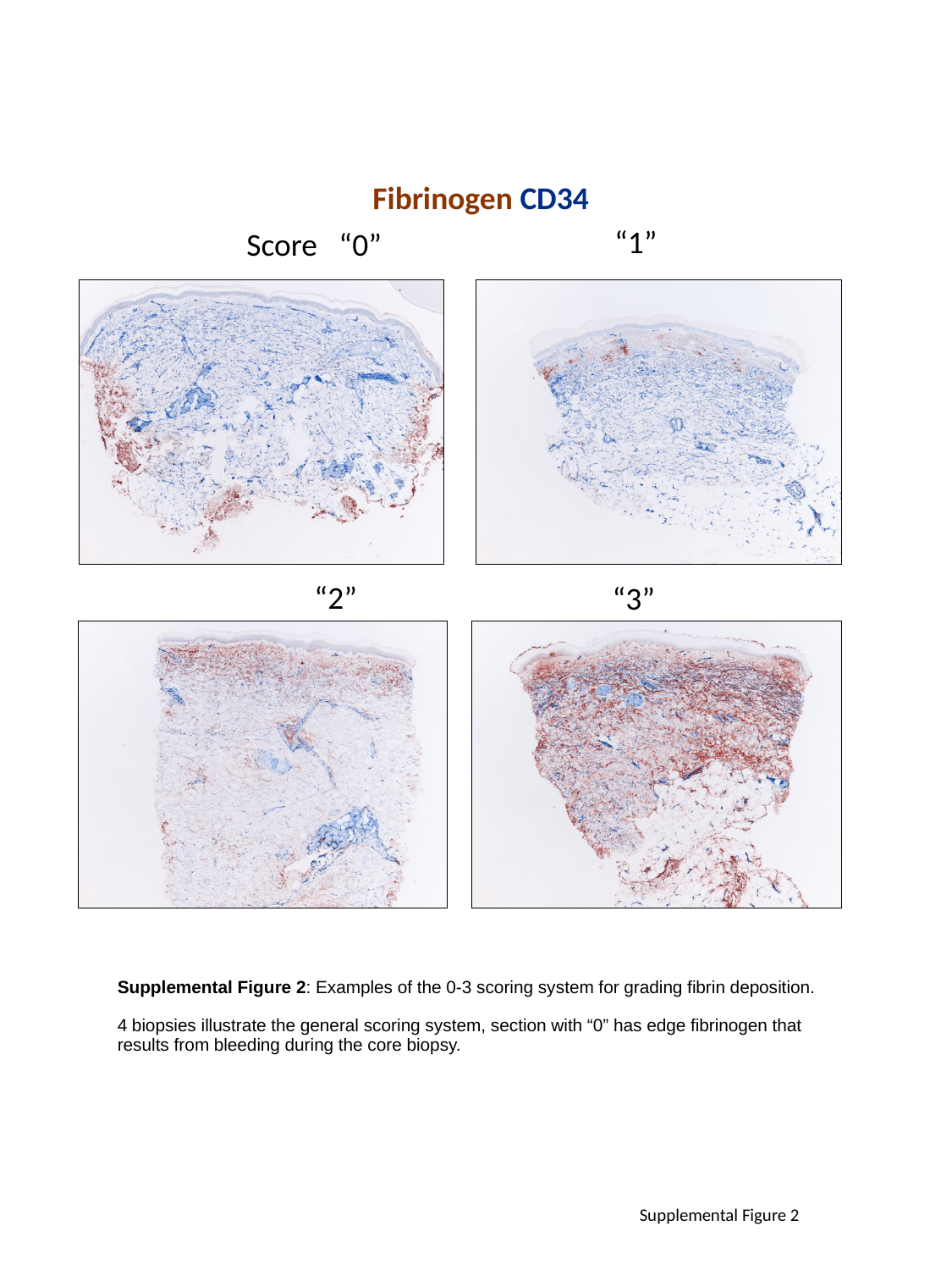

Fibrinogen CD34
“1”
Score “0”
“2”
“3”
#### Supplemental Figure 2: Examples of the 0-3 scoring system for grading fibrin deposition. 4 biopsies illustrate the general scoring system, section with “0” has edge fibrinogen that results from bleeding during the core biopsy.
Supplemental Figure 2

### Slide 5
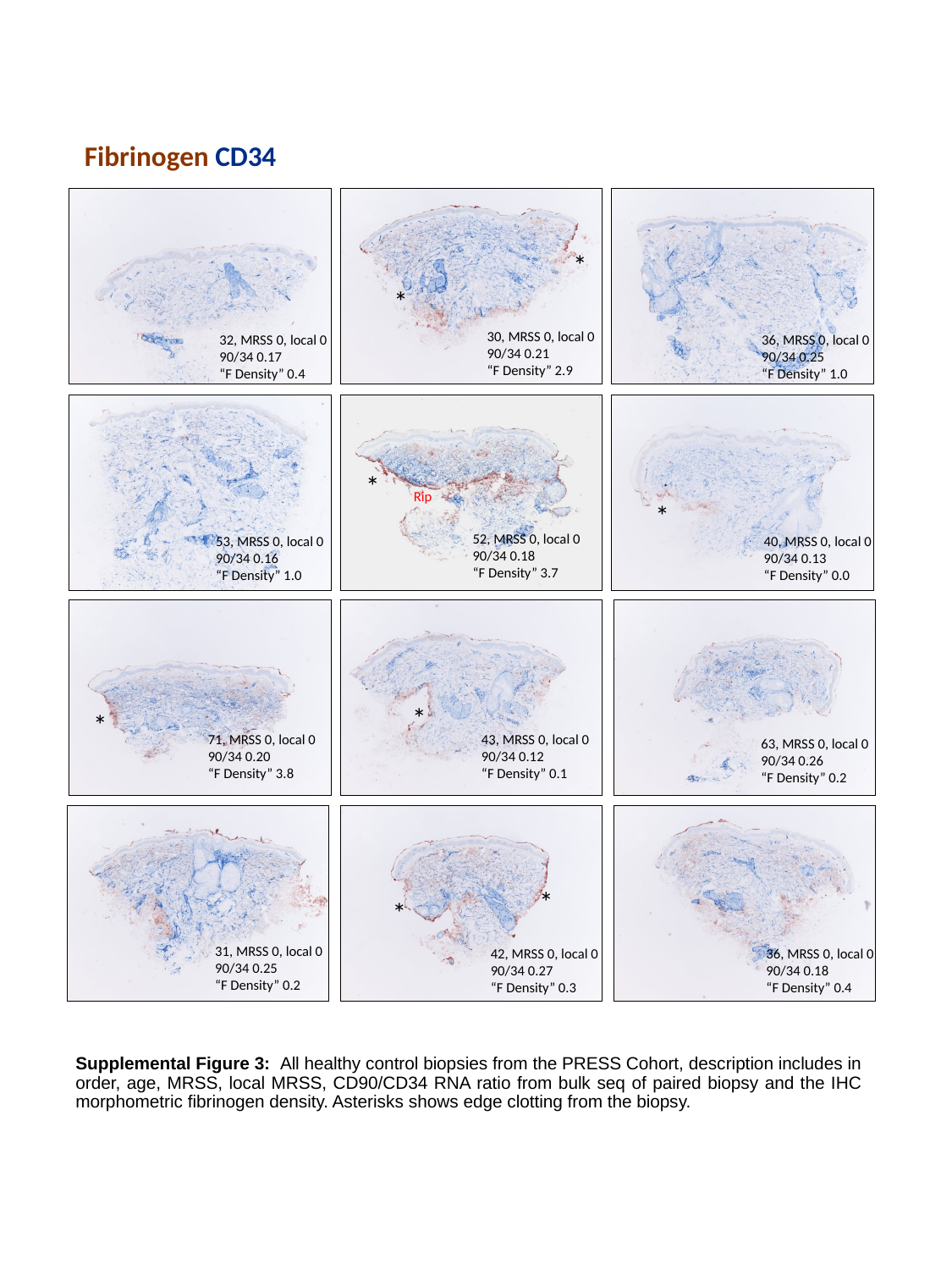

Fibrinogen CD34
*
*
30, MRSS 0, local 0
90/34 0.21
“F Density” 2.9
36, MRSS 0, local 0
90/34 0.25
“F Density” 1.0
32, MRSS 0, local 0
90/34 0.17
“F Density” 0.4
*
Rip
*
52, MRSS 0, local 0
90/34 0.18
“F Density” 3.7
53, MRSS 0, local 0
90/34 0.16
“F Density” 1.0
40, MRSS 0, local 0
90/34 0.13
“F Density” 0.0
*
*
43, MRSS 0, local 0
90/34 0.12
“F Density” 0.1
71, MRSS 0, local 0
90/34 0.20
“F Density” 3.8
63, MRSS 0, local 0
90/34 0.26
“F Density” 0.2
*
*
31, MRSS 0, local 0
90/34 0.25
“F Density” 0.2
42, MRSS 0, local 0
90/34 0.27
“F Density” 0.3
36, MRSS 0, local 0
90/34 0.18
“F Density” 0.4
#### Supplemental Figure 3: All healthy control biopsies from the PRESS Cohort, description includes in order, age, MRSS, local MRSS, CD90/CD34 RNA ratio from bulk seq of paired biopsy and the IHC morphometric fibrinogen density. Asterisks shows edge clotting from the biopsy.

### Slide 6
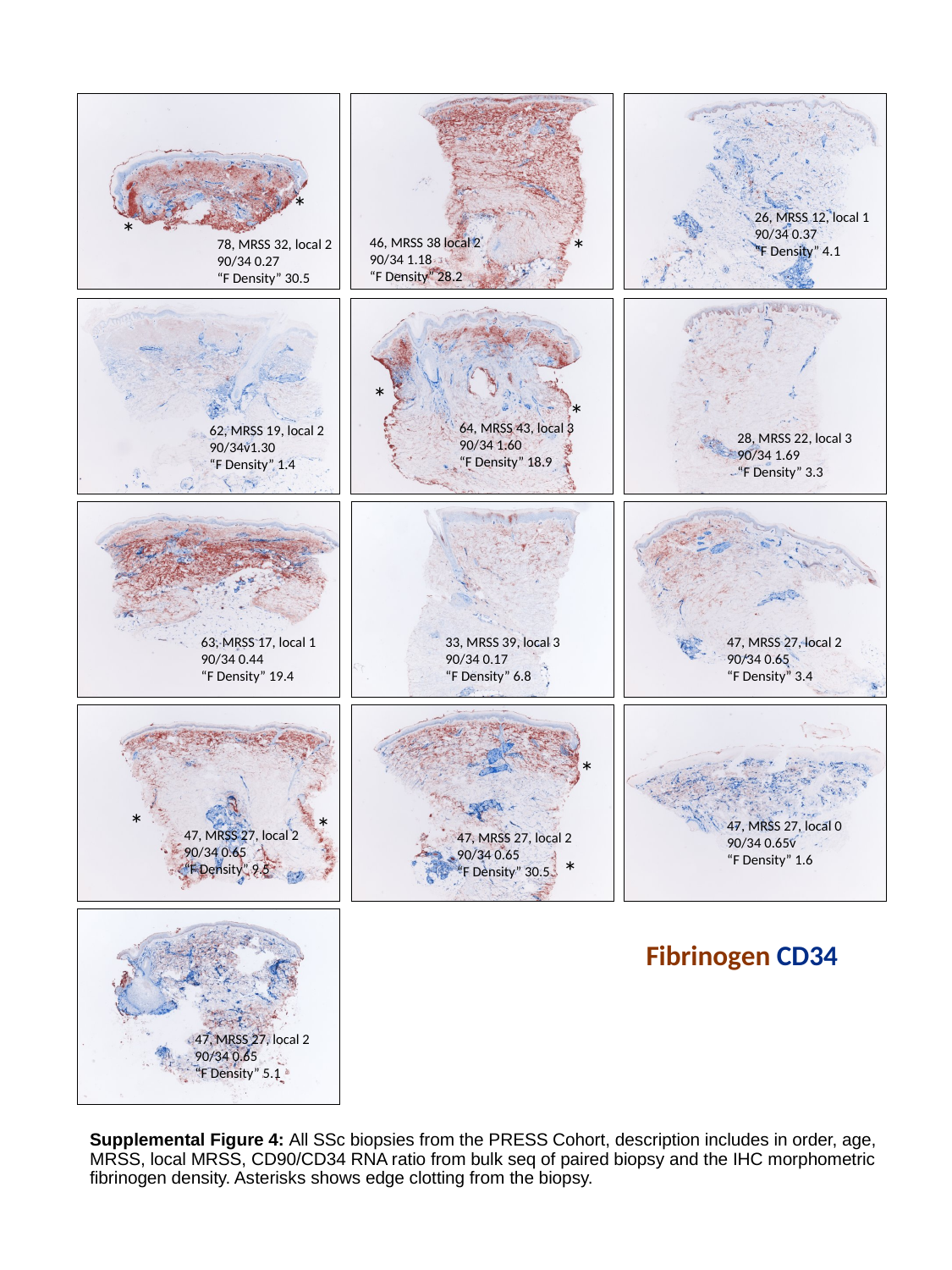

*
26, MRSS 12, local 1
90/34 0.37
“F Density” 4.1
*
*
46, MRSS 38 local 2
90/34 1.18
“F Density” 28.2
78, MRSS 32, local 2
90/34 0.27
“F Density” 30.5
*
*
64, MRSS 43, local 3
90/34 1.60
“F Density” 18.9
62, MRSS 19, local 2
90/34v1.30
“F Density” 1.4
28, MRSS 22, local 3
90/34 1.69
“F Density” 3.3
63, MRSS 17, local 1
90/34 0.44
“F Density” 19.4
33, MRSS 39, local 3
90/34 0.17
“F Density” 6.8
47, MRSS 27, local 2
90/34 0.65
“F Density” 3.4
*
*
*
47, MRSS 27, local 0
90/34 0.65v
“F Density” 1.6
47, MRSS 27, local 2
90/34 0.65
“F Density” 9.5
47, MRSS 27, local 2
90/34 0.65
“F Density” 30.5
*
Fibrinogen CD34
47, MRSS 27, local 2
90/34 0.65
“F Density” 5.1
Supplemental Figure 4: All SSc biopsies from the PRESS Cohort, description includes in order, age, MRSS, local MRSS, CD90/CD34 RNA ratio from bulk seq of paired biopsy and the IHC morphometric fibrinogen density. Asterisks shows edge clotting from the biopsy.

### Slide 7
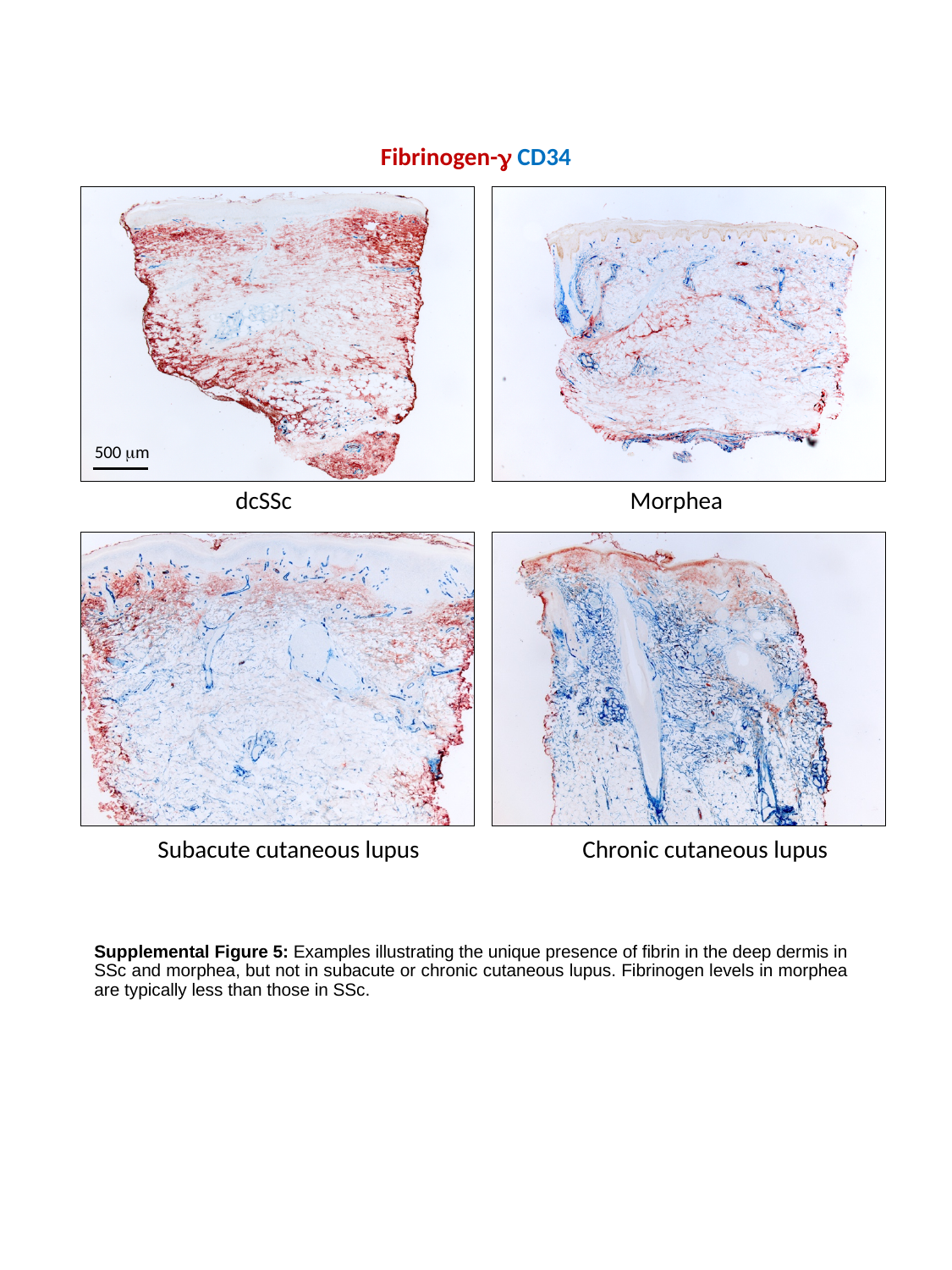

Fibrinogen-γ CD34
500 μm
dcSSc
Morphea
Subacute cutaneous lupus
Chronic cutaneous lupus
Supplemental Figure 5: Examples illustrating the unique presence of fibrin in the deep dermis in SSc and morphea, but not in subacute or chronic cutaneous lupus. Fibrinogen levels in morphea are typically less than those in SSc.

### Slide 8
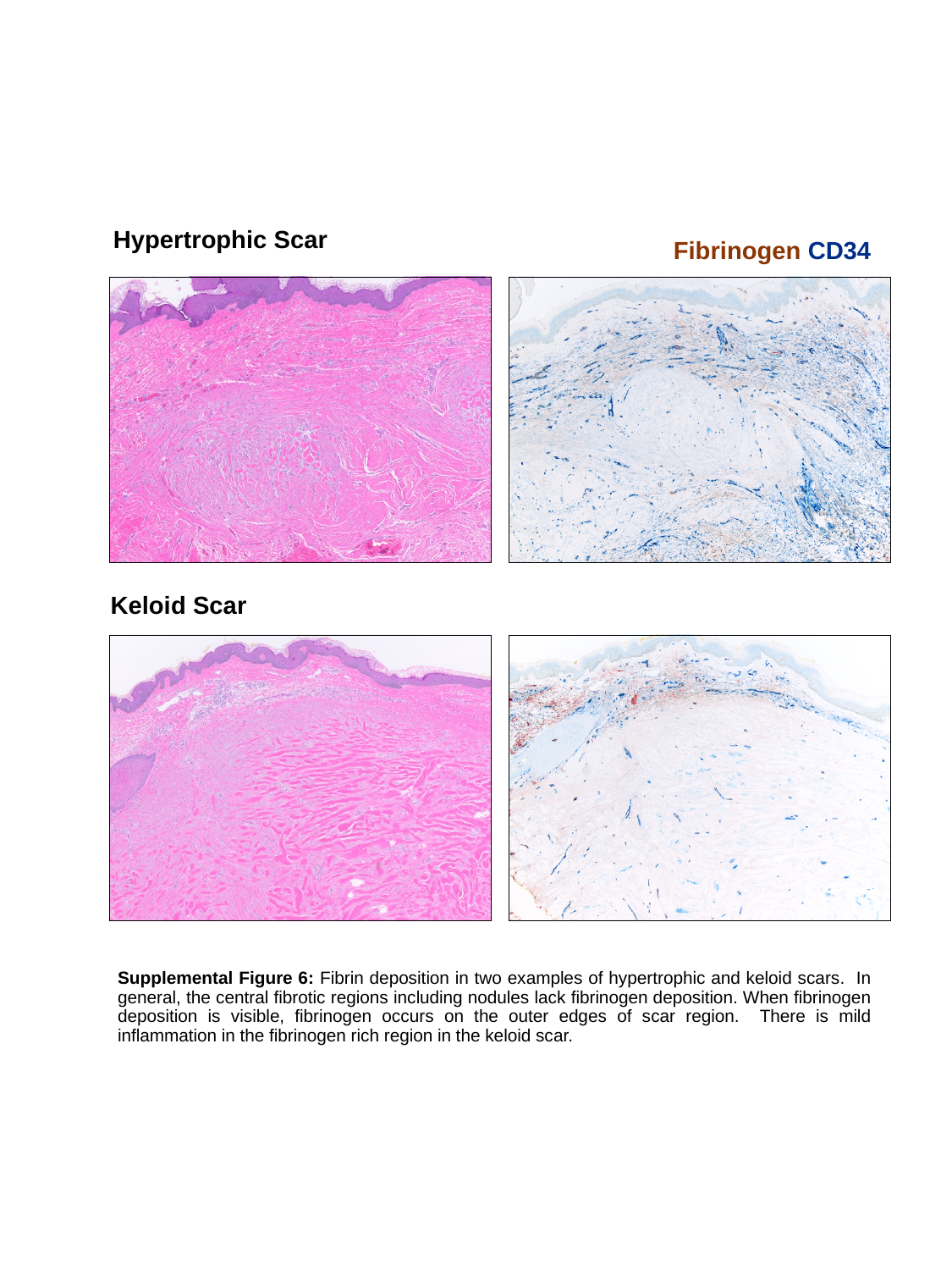

Hypertrophic Scar
Fibrinogen CD34
Keloid Scar
Supplemental Figure 6: Fibrin deposition in two examples of hypertrophic and keloid scars. In general, the central fibrotic regions including nodules lack fibrinogen deposition. When fibrinogen deposition is visible, fibrinogen occurs on the outer edges of scar region. There is mild inflammation in the fibrinogen rich region in the keloid scar.

### Slide 9
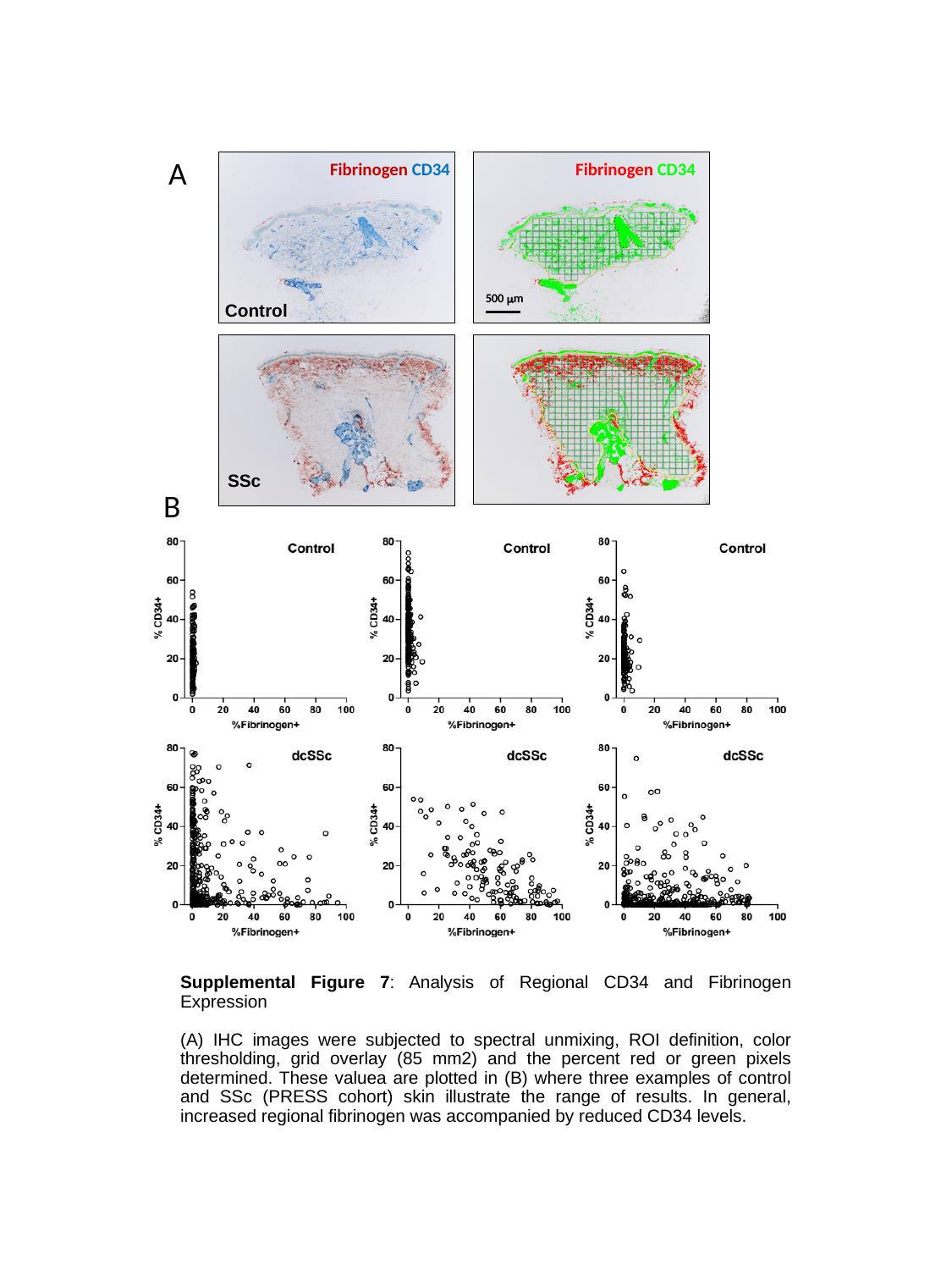

A
Fibrinogen CD34
Fibrinogen CD34
Control
SSc
500 μm
B
Supplemental Figure 7: Analysis of Regional CD34 and Fibrinogen Expression
(A) IHC images were subjected to spectral unmixing, ROI definition, color thresholding, grid overlay (85 mm2) and the percent red or green pixels determined. These valuea are plotted in (B) where three examples of control and SSc (PRESS cohort) skin illustrate the range of results. In general, increased regional fibrinogen was accompanied by reduced CD34 levels.

### Slide 10
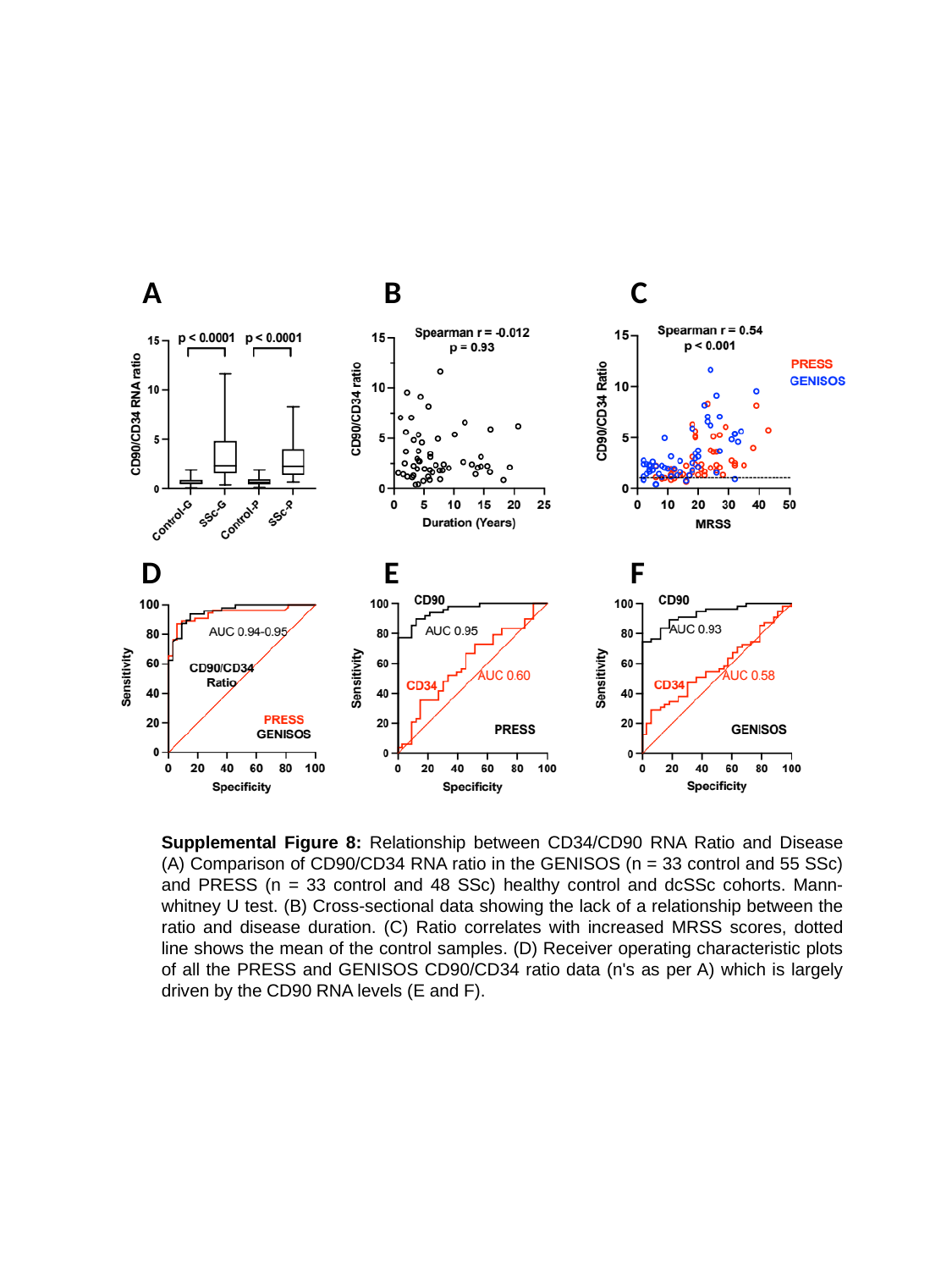

A
B
C
D
E
F
Supplemental Figure 8: Relationship between CD34/CD90 RNA Ratio and Disease (A) Comparison of CD90/CD34 RNA ratio in the GENISOS (n = 33 control and 55 SSc) and PRESS (n = 33 control and 48 SSc) healthy control and dcSSc cohorts. Mann-whitney U test. (B) Cross-sectional data showing the lack of a relationship between the ratio and disease duration. (C) Ratio correlates with increased MRSS scores, dotted line shows the mean of the control samples. (D) Receiver operating characteristic plots of all the PRESS and GENISOS CD90/CD34 ratio data (n's as per A) which is largely driven by the CD90 RNA levels (E and F).
